# Alternatively polarized neutrophils metabolically restrain macrophage-driven inflammation

**DOI:** 10.64898/2026.09.30.755643

**Authors:** Paul Ettel, Jayne Louise Wilson, Piyal Saha, Mario Mazic, Aglaja Kopf, Kveta Brazdilova, Georg Stary, Ivan Melikhov, Anne Miller, Jadwiga Jablonska, Markus H Hoffmann, Gernot Schabbauer, Markus Hengstschläger, Thomas Weichhart

## Abstract

Macrophage-driven inflammation underlies many inflammatory diseases, yet the mechanisms that restrain macrophage activation within inflamed tissues remain poorly understood. Using mouse models of foreign body granuloma formation and macrophage activation syndrome, we show that neutrophils accumulate at sites of inflammation and metabolically restrain macrophage-driven pathology. Neutrophil depletion or genetic impairment exacerbated tissue pathology, whereas induced neutrophilia attenuated inflammatory damage. Tissue-infiltrating neutrophils acquired an alternatively polarized state characterized by high arginase 1 activity that was regulated by an IL4Rα–STAT3–NOX2 signaling axis. Functionally, polarized neutrophils depleted extracellular arginine in an arginase 1-dependent manner, thereby limiting macrophage effector programs. Human neutrophils exhibited conserved regulation of arginase activity, and alternatively polarized neutrophils were detected in patients with sarcoidosis and macrophage activation syndrome. Together, these findings identify an arginine-dependent metabolic mechanism through which alternatively polarized neutrophils restrain macrophage-driven inflammation and reveal a previously unrecognized immune-regulatory role for these cells in chronic inflammatory disease.

## Introduction

Inflammatory circuits are often modeled as self-amplifying cascades where myeloid cells reinforce pathological tissue responses, yet counter-regulatory mechanisms that impose brakes on chronic inflammation remain poorly defined^1,2^. Macrophages are central innate immune regulators of tissue inflammation and repair. While they are essential for tissue homeostasis, they also represent early responders to danger signals and potent initiators of pathological inflammation^1^. In chronic inflammatory diseases such as granulomatous disorders and macrophage activation syndrome (MAS), sustained macrophage activation drives pathological tissue remodeling and damage. Granulomas are dense, macrophage-rich structures that form in response to foreign bodies (foreign body granuloma), infectious agents (*e.g.*, *Mycobacterium tuberculosis*), or autoinflammatory diseases like sarcoidosis^3,4^. Although the triggers for granuloma formation differ, macrophages usually aggregate, subsequently undergo epithelioid differentiation and can develop into multinucleated giant cells (MGCs)^3,5^. In MAS, which occurs in patients with autoinflammatory diseases or cancer, dysregulated macrophages trigger pancytopenia and tissue damage through excessive phagocytosis and cytokine secretion^6^. While macrophage-intrinsic signaling and metabolic pathways promoting these responses are increasingly understood^4,7,8^, the intercellular mechanisms restraining macrophage-driven inflammation remain incompletely defined.

Neutrophils, the most abundant circulating leukocytes in humans, are rapidly recruited from bone marrow to sites of infection and tissue injury^9^. While historically considered short-lived effector cells dedicated to clearing pathogens^10,11^, neutrophils are increasingly recognized as important regulators of tissue homeostasis^12,13^. In sterile inflammation, neutrophil accumulation is widely viewed as a pro-inflammatory stimulus that can lead to tissue damage. Yet, in situations such as surgical trauma or prolonged metabolic stress, this classic view does not fully explain their contribution to disease^14^. Neutrophils can shape macrophage activation states through efferocytosis and tissue-derived signals^15–18^ and impaired clearance of neutrophils can lead to prolonged tissue inflammation^19,20^. Although neutrophils are considered early recruiters of monocytes to sites of granulomatous inflammation^21,22^, whether they directly regulate macrophage effector programs during chronic inflammation remains unknown. Recently, cytokine-induced alternatively polarized neutrophils were identified as a distinct, immature Ly6G^low^CD14^high^CD101^low^ subset that promotes neuronal regeneration and axon regrowth following CNS injury^23,24^. While this subset exhibits pro-regenerative features in neuronal contexts, their broader existence, their role in regulating chronic inflammatory disease, as well as the signaling pathways controlling their polarization remain poorly understood.

Here, using complementary murine models of foreign body granuloma formation, *Tsc2*-deficient sarcoidosis-like disease, and CpG-induced MAS, we show that neutrophils rapidly accumulated at sites of inflammation to metabolically restrain macrophage-driven pathology. Neutrophil depletion or genetic impairment (*e.g.,* NOX2 deficiency) exacerbated granulomatous inflammation and organ damage, whereas induced neutrophilia attenuated disease. Tissue-infiltrating neutrophils acquired an alternatively polarized, Arginase 1 (Arg1)-high state that depended on an IL4Rα-STAT3-NOX2 signaling axis. Functionally, these alternatively polarized neutrophils depleted extracellular arginine via Arg1 to limit macrophage effector programs and reduced granulomatous inflammation *in vivo.* Moreover, neutrophils were detected in sarcoidosis and MAS patients, and human neutrophils exhibited conserved Arg1 regulation and granuloma-modulating capacity *in vitro*. Together, these findings identify an alternatively polarized neutrophil state that metabolically restrains macrophage-driven inflammation in chronic inflammatory disease.

## Results

### Neutrophils restrain macrophage-driven inflammatory pathology

As intercellular mechanisms that restrain macrophage-driven inflammation remain poorly defined, we investigated whether neutrophils regulate this process *in vivo*. We initially employed a foreign body granuloma model^25–28^, in which granuloma formation is initiated by subcutaneous implantation of glass coverslips in the rear flanks of mice (**Figure 1A**). This led to the infiltration of neutrophils and monocytes, that further developed into macrophages and ultimately formed MGCs by day 9 (**Figure 1B**). Neutrophils dominated the early inflammatory infiltrate but were also observed at later stages of granuloma maturation (**Figure 1C, Figure S1A, B**). Granuloma-associated neutrophils showed an activated phenotype indicated by reduced Ly6G and CD62L expression and increased CD11b levels compared to circulating neutrophils (**Figure S1C-E**). To determine whether they functionally influence granuloma formation, we depleted neutrophils using an Ly6G antibody prior to coverslip implantation^29^ (**Figure 1D, Figure S1F, G**). Surprisingly^21,22^, neutrophil depletion markedly increased granuloma development nine days after implantation (**Figure 1E, F**). Similar effects were observed when neutrophils were depleted for longer periods, suggesting that neutrophils restrain rather than promote granulomatous inflammation (**Figure S1G-J**). To genetically validate these findings, we implanted mice functionally deficient in the NADPH oxidase component NOX2 (denoted *Ncf1\*\**), a model of chronic granulomatous disease characterized by defective neutrophil reactive oxygen species (ROS) production^30,31^ (**Figure 1G**)^32–34^. Like neutrophil-depleted animals, *Ncf1\*\** mice displayed exacerbated granuloma formation, indicating that intact neutrophil effector functions are required to restrain granulomatous inflammation (**Figure 1H, I**). We then asked whether increasing neutrophil abundance (neutrophilia) would suppress granuloma development. Indeed, administration of granulocyte-colony stimulating factor (G-CSF) (**Figure 1J**)^29^ boosted circulating neutrophil numbers (**Figure S1K, L**) and markedly restrained granulomatous inflammation *in vivo* (**Figure 1K, L**).

**Figure 1:**
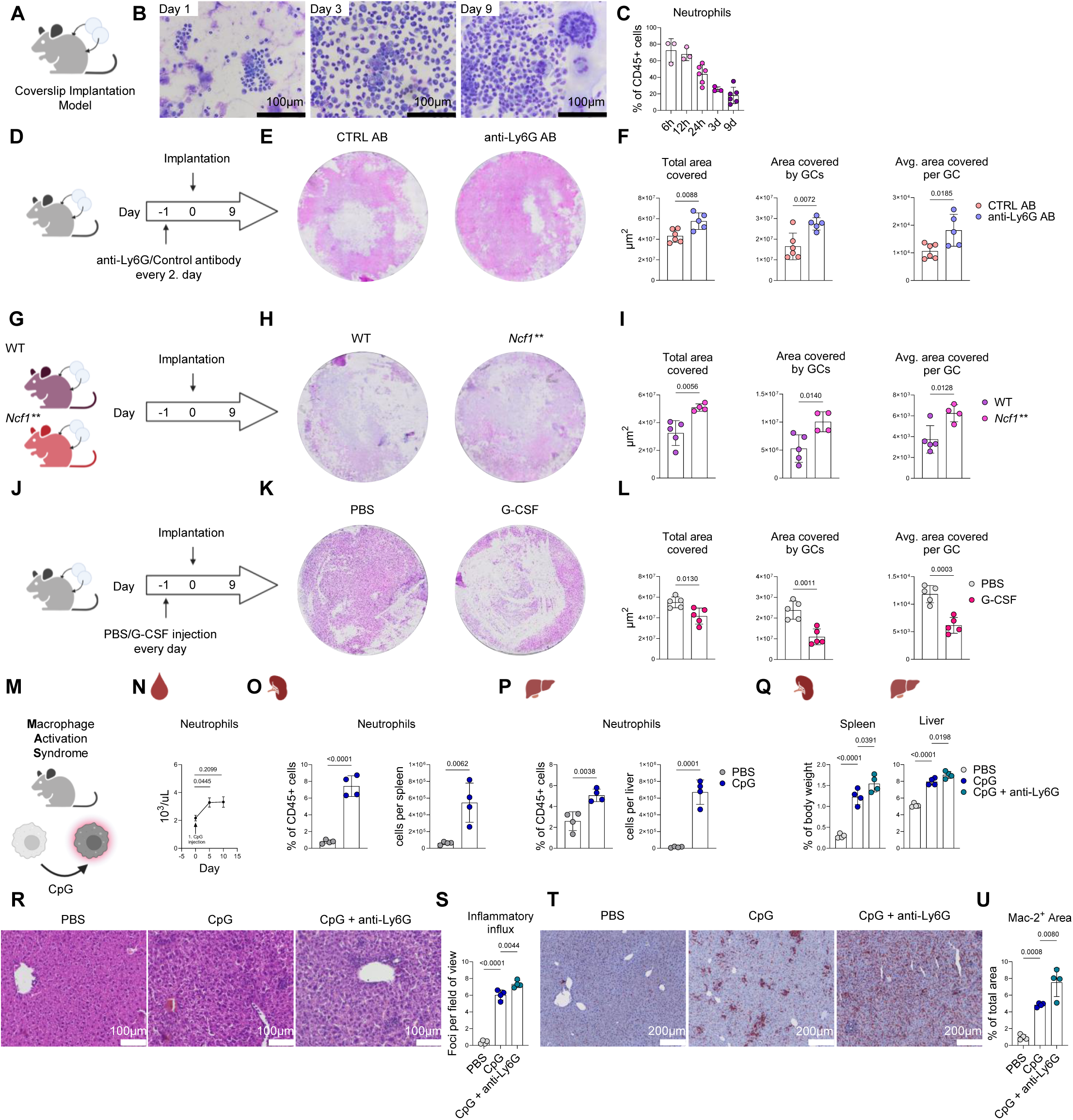
Neutrophils limit macrophage-driven inflammation *in vivo*. **(A)** Schematic of the coverslip implantation model of granuloma development. **(B)** Representative May-Grünwald-Giemsa staining of cells attached to the coverslip 1, 3, and 9 days after implantation. **(C)** Frequency of neutrophils among CD45+ cells at indicated time points (n = 3 - 6 per time point). **(D)** Experimental design for neutrophil depletion using anti-Ly6G during granuloma formation. **(E)** Representative H&E staining of coverslips from control or anti-Ly6G-depleted mice at day 9. **(F)** Quantification of granuloma burden by total area covered, giant cell area (GCs), and average (Avg.) GC size (n = 5 - 6 per group). **(G)** Experimental design for coverslip implantation in wild-type (WT) and *Ncf1\*\** mice. **(H)** Representative H&E staining of coverslips in WT and *Ncf1\*\** mice. **(I)** Quantification of granuloma burden in WT or *Ncf1\*\** mice (n = 4 – 5 per genotype). **(J)** Experimental design of coverslip implantation in mice receiving PBS or G-CSF injections. **(K)** Representative H&E staining of coverslips from PBS- or G-CSF-treated mice. **(L)** Quantification of granuloma burden 9 days after CS implantation in mice receiving PBS or G-CSF (n = 5 per group). **(M)** Schematic of the CpG-induced macrophage activation syndrome (MAS) model. **(N)** Neutrophil counts in peripheral blood at indicated time points after the first CpG administration (n = 4). **(O, P)** Neutrophil frequency and absolute numbers in spleen (**O**) and liver (**P**) following PBS or CpG treatment (n = 4 per group). **(Q)** Splenic and hepatic weight as percentage of total body weight in mice upon PBS, CpG, or CpG and anti-Ly6G treatment (n = 4 per group). **(R)** Representative H&E staining and **(S)** quantification of inflammatory influx of liver samples. **(T)** Representative Mac2 staining by IHC and **(U)** quantification. **(C, F, I, L, O, P, Q, S, U)** Data are shown as means ± SD and **(N)** mean ± SEM. Statistical analysis: unpaired t test **(F, I, L, O, P);** RM one-way ANOVA (**N**); one-way ANOVA **(Q, S, U)**.

To investigate whether neutrophils limit macrophage-driven inflammation in other contexts, we used a mouse model of MAS induced by injecting the TLR9 agonist CpG^7,35,36^(**Figure 1M**). CpG administration led to a rapid and sustained increase in neutrophil numbers in peripheral blood (**Figure 1N**) and a pronounced accumulation of neutrophils in the spleen and liver, two organs prone to tissue damage in this model (**Figure 1O, P; Figure S2A, B**). To determine their functional relevance in this context, we depleted neutrophils with an Ly6G antibody as described above (**Figure S2C**). Neutrophil depletion significantly exacerbated disease severity indicated by increased splenomegaly and hepatomegaly (**Figure 1Q**). Further histological analysis revealed an increase in the accumulation in inflammatory foci, a hallmark of the model, in the livers of neutrophil-depleted mice (**Figure 1R, S**). Consistent with exacerbated disease pathology, neutrophil depletion led to a further expansion of macrophages as compared to CpG-treated mice alone (**Figure 1T, U**). These findings demonstrate that neutrophils restrain macrophage-driven pathology across distinct inflammatory contexts, prompting us to investigate the mechanisms underlying this regulatory function.

### Neutrophils acquire an alternative polarization state in macrophage-driven inflammation

To define the molecular programs associated with the inflammation-limiting phenotype, we performed bulk RNA sequencing on FACS-sorted neutrophils from the blood or coverslip three days after implantation (**Figure 2A**). Principle component analysis (PCA) revealed clear transcriptional differences between circulating and granuloma-infiltrating neutrophils (**Figure 2B**), with thousands of differentially expressed genes identified between the groups (**Figure S3A**). Granuloma neutrophils displayed features of classical activation, including increased expression of genes involved in neutrophil adhesion (*Icam1*), neutrophil swarming (*Tnf*), and cellular lifespan (*Fas*), consistent with the activated phenotype observed by flow cytometry (**Figure 2C, S1BC-E**). Additionally, gene ontology analysis showed enrichment of inflammatory signaling pathways including NF-κB and interferon signaling (**Figure S3B**). Given the granuloma-limiting phenotype of neutrophils, we speculated that they might acquire a suppressive phenotype, similar to those described for myeloid-derived suppressor cells (MDSC)^37^. However, evaluation of MDSC markers^38^ showed only a partial overlap with the granuloma neutrophils (**Figure 2D**). Instead, the transcriptional profile closely resembled recently described signatures of alternatively polarized neutrophils (**Figure 2E**)^24^. Flow cytometric analysis confirmed the expression of key markers for alternative polarization like Ly6G, F4/80, the maturation marker CD101, and enzymes involved in arginine metabolism such as inducible nitric oxide synthase (iNOS) and arginase 1 (Arg1) (**Figure 2F - J**). Notably, hepatic neutrophils in the MAS model displayed a comparable polarization phenotype, indicating that alternative polarization represents a conserved neutrophil state in macrophage-driven inflammation (**Figure 2K, L**).

**Figure 2:**
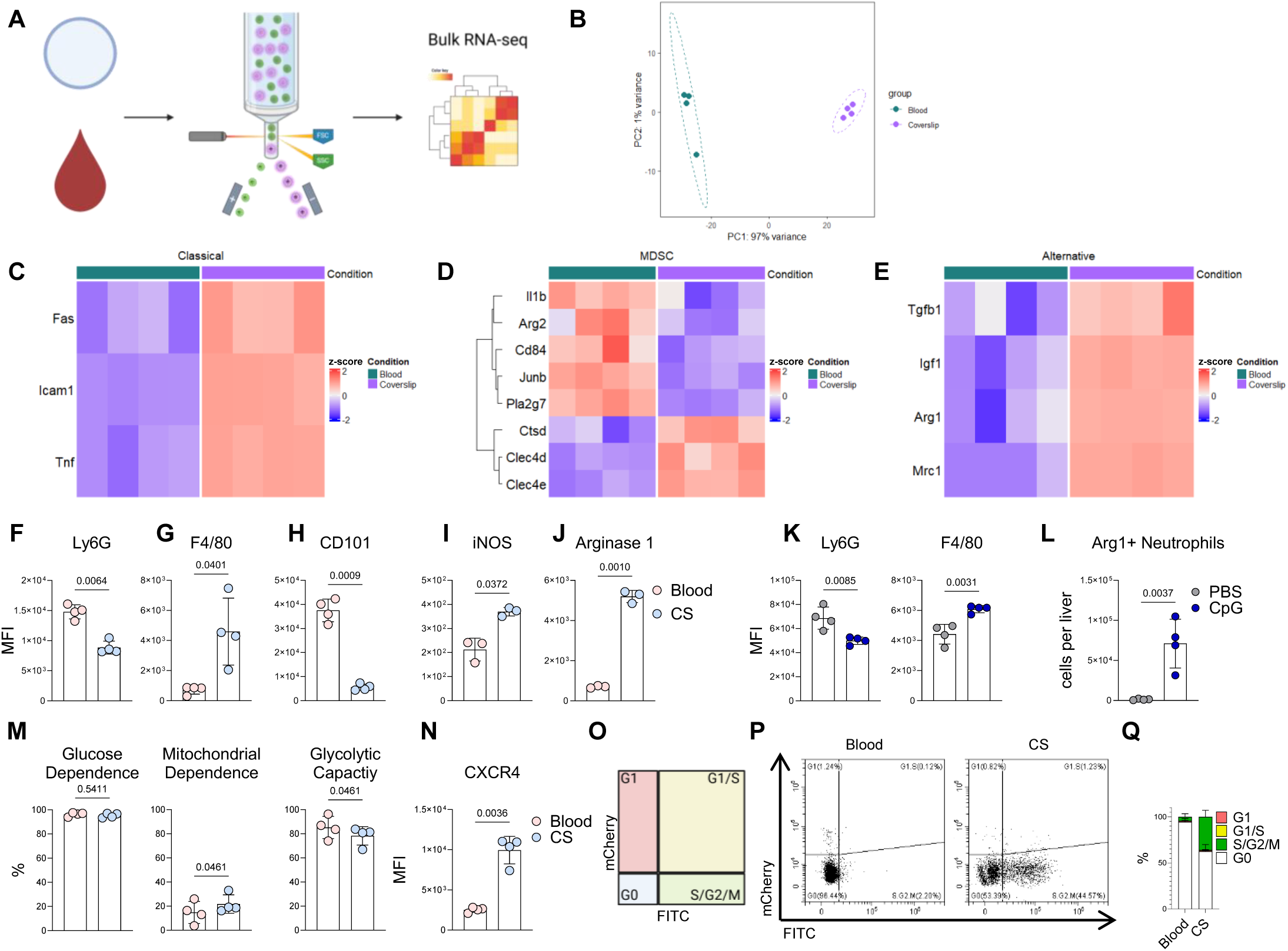
RNA sequencing reveals alternative polarization of neutrophils in macrophage-driven inflammation. **(A)** Experimental design for bulk RNA sequencing of FACS-sorted neutrophils from blood and coverslips. **(B)** Principal component analysis (PCA) plot of neutrophils transcriptomes as outlined in **(A)**. Heat maps of signatures for classical neutrophil activation **(C)**, myeloid-derived suppressor cells (MDSC**; D)**, and alternative neutrophil polarization **(E).** Flow cytometry quantification of **(F)** Ly6G, **(G)** F4/80, **(H)** CD101, **(I)** iNOS, and **(J)** Arg1 in neutrophils from blood and coverslip 3 days after implantation (n = 3 - 4). **(K, L)** Analysis of neutrophil polarization in the MAS model. **(K)** Expression of Ly6G (left) and F4/80 (right) in hepatic neutrophils from PBS- or CpG-treated mice (n = 4 per condition). **(L)** Quantification of Arg1 + neutrophils per liver. **(M)** Metabolic profiling of glucose dependence, mitochondrial dependence, and glycolytic capacity in neutrophils isolated from blood and CS 3 days after implantation using SCENITH^TM^ (n = 4). (**N)** Expression of CXCR4 on neutrophils 3 days after implantation (n = 4). **(O)** Schematic of cell cycle analysis using Fucci reporter mice. **(P)** Representative flow cytometry plots of neutrophils from blood and coverslips using Fucci mice 3 days after implantation. **(Q)** Quantification of cell cycle phases of neutrophils in Fucci mice 3 days after implantation (n = 4). Data are shown as means ± SD. Statistical analyses: paired student t tests **(F-J, M, N),** unpaired student t test (**K-L**).

Metabolically, the granuloma neutrophils stayed strongly dependent on glycolysis, similar to circulating neutrophils but showed a higher mitochondrial dependence (**Figure 2M**). Markers associated with immature neutrophil populations, including reduced CD101 expression and increased CXCR4 levels^39^, were also detected (**Figure 2H, N**). Indeed, using transgenic Fucci cell cycle reporter mice^40^ a proliferative capacity of granuloma neutrophils in contrast to postmitotic neutrophils in blood was observed (**Figure 2O – Q**). Collectively, we observed that neutrophils in models of macrophage-driven inflammation undergo extensive transcriptional reprogramming and acquire an alternative polarized phenotype characterized by Arg1 expression, suggesting that these cells may regulate macrophage responses through arginine metabolism.

### Alternative neutrophil polarization is controlled by an IL4Rα-STAT3-NOX2 axis

Having identified alternatively polarized neutrophils in macrophage-driven inflammation, we next sought to define the signaling pathways that regulate this state. We therefore analyzed enriched pathways in the RNA sequencing data set, which revealed IL6-JAK-STAT3 signaling as one of the most enriched ones (**Figure S4A, B**). Consistently, neutrophils isolated from coverslip granulomas displayed increased phosphorylation of STAT3 compared to circulating neutrophils (**Figure 3A, B**). Given the emerging evidence on STAT3 signaling in controlling neutrophil effector functions^41,42^, we investigated whether STAT3 signaling controls alternative polarization. Generation of alternatively polarized bone marrow-derived neutrophils (Pol BMNs) via stimulation with IL4 and G-CSF (**Figure 3C**)^24^ led to a potent increase in STAT3 phosphorylation *in vitro* that stabilized after 1h (**Figure 3D**). Moreover, IL4 and G-CSF stimulation resulted in strong induction of Arg1, which is one of the most strongly upregulated markers of alternatively polarized BMNs. Pharmacological inhibition of STAT3 (STAT3i) markedly dampened the expression of Arg1 (**Figure 3E, F**) and partially blocked additional markers of alternative polarization (**Figure S4C**).

**Figure 3:**
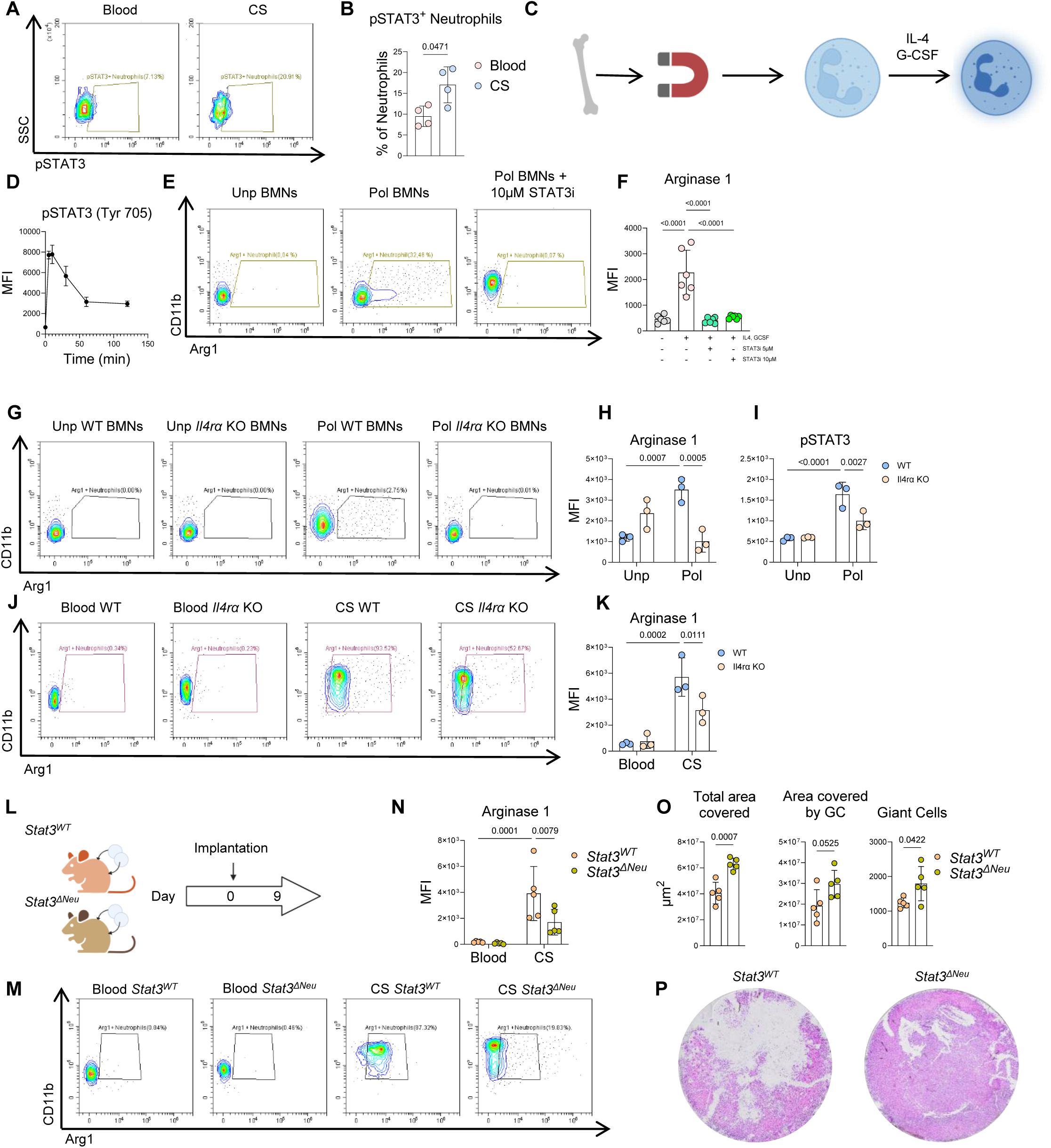
An IL4Ra-STAT3-NOX2 signaling axis drives Arginase 1 expression and neutrophil-mediated macrophage inflammation. **(A)** Representative flow cytometry plots and quantification **(B)** of pSTAT3+ neutrophils in blood and CS 3 days after implantation (n = 4). **(C)** Experimental design for *in vitro* generation of alternatively polarized bone marrow derived-neutrophils (Pol BMNs). **(D)** Kinetics of STAT3 phosphorylation in neutrophils stimulated with IL4 and G-CSF (n = 3). **(E)** Representative flow cytometry plots and **(F)** quantification of arginase 1 expression 24h after stimulation in unpolarized (Unp) BMNs and Pol BMNs in the presence of absence of a STAT3 inhibitor (STAT3i) (n = 6). **(G)** Representative flow cytometry plots and **(H)** quantification of arginase 1 expression in WT and *Il4ra* KO neutrophils left unpolarized or in the presence of IL4 and G-CSF (n = 3 per genotype). **(I)** STAT3 phosphorylation in WT and *Il4ra* KO US neutrophils and Pol BMNs for 24h (n = 3 per genotype). **(J)** Representative flow cytometry plots and **(K)** quantification of arginase 1 expression in blood and coverslips (CS) from WT and *Il4ra* KO neutrophils 3d after implantation (n = 3 per genotype). **(L)** Experimental design for coverslip implantation in *Stat3* WT (*Stat3^WT^*) and neutrophil specific *Stat3* deficient (*Stat3^ΔNeu^*) mice. **(M)** Representative flow cytometry plots and **(N)** quantification of Arg1 in neutrophils in blood and cover slips of *Stat3^WT^* and *Stat3^ΔNeu^* mice 9 days after CS implantation. **(O)** Quantification and **(P)** representative images of granuloma burden in *Stat3^W^*^T^ and *Stat3^ΔNe^*^u^ mice at the experimental endpoint (n = 5 per genotype). Data are shown as means ± SD **(B, D, F, H, I K, N, O)**. Statistical analysis: paired student t test **(B);** one-way ANOVA **(F);**2-way ANOVA **(H, I, K, N);** unpaired student t test **(O)**.

Given the stimulation with IL4, we hypothesized that the IL4 receptor (IL4Rα) might regulate polarization upstream of STAT3. Neutrophils isolated from *Il4rα*-deficient mice showed reduced Arg1 and F4/80 expression upon cytokine stimulation (**Figure 3G, H, Figure S4D**) and exhibited impaired STAT3 phosphorylation (**Figure 3I**). Consistent with these findings, *Il4rα*-deficient mice reduced Arg1 expression in granuloma neutrophils in the coverslip implantation model *in vivo* (**Figure 3J, K**).

Based on the observation that NOX2 deficiency in the *Ncf1\*\** mice exacerbated granuloma formation *in vivo* (**Figure 1H, I**), we next investigated whether ROS contribute to neutrophil polarization. Pharmacological inhibition of NOX2 strongly suppressed Arg1 induction (**Figure S4E, F**), while other markers were unaffected (**Figure S4G**). Similarly, directly scavenging ROS using N-acetyl cysteine (NAC) reduced Arg1 expression (**Figure S4H**). These findings indicate that NOX2-dependent ROS selectively regulate Arg1 expression, linking classical neutrophil effector functions to the metabolic program that restrains macrophage-driven inflammation. Evaluation of a potential crosstalk between STAT3 and NOX2 showed that STAT3 inhibition reduced NOX2 expression (**Figure S4I**), whereas NOX2 inhibition did not affect STAT3 activation, suggesting that NOX2 functions downstream of STAT3 (**Figure S4J)**. Together, these findings identify an Il4Rα-STAT3-NOX2 signaling axis regulating Arg1 expression during neutrophil polarization.

To determine whether this pathway is functionally relevant *in vivo*, we analyzed neutrophil-specific *Stat3*-deficient mice (*Stat3^ΔNeu^*) using the coverslip granuloma model (**Figure 3L**)^42^. *Stat3*-deficient granuloma neutrophils showed reduced expression of Arg1 (**Figure 3M, N**), F4/80, and CXCR4 (**Figure S4K**) *in vivo*, while other polarization markers remained largely unchanged (**Figure S4K**). Importantly, *Stat3^ΔNe^*^u^ mice showed significantly exacerbated granuloma formation (**Figure 3O, P**), demonstrating that STAT3-dependent neutrophil function restrains macrophage-driven inflammation *in vivo*.

### Neutrophils restrict macrophage effector functions by limiting arginine availability

Arginine metabolism is a key regulator of macrophage effector functions and MGC formation^43^. As coverslip macrophages highly expressed Arg1 (**Figure S5A**), we examined whether arginine availability controls granuloma formation using a macrophage clustering model *in vitro*. Bone marrow-derived macrophages (BMDMs) from myeloid-specific *Tsc2*-deficient mice, which develop spontaneous sarcoid-like granulomas *in vivo,* form granuloma-like clusters *in vitro*^44,45^. Removal of arginine from the culture medium strongly abrogated cluster formation in this *in vitro* model (**Figure 4A, B**). Conversely, increasing extracellular arginine concentrations enhanced macrophage clustering in a dose-dependent manner (**Figure 4C**). These findings indicate that extracellular arginine availability directly regulates macrophage-driven granuloma formation.

**Figure 4:**
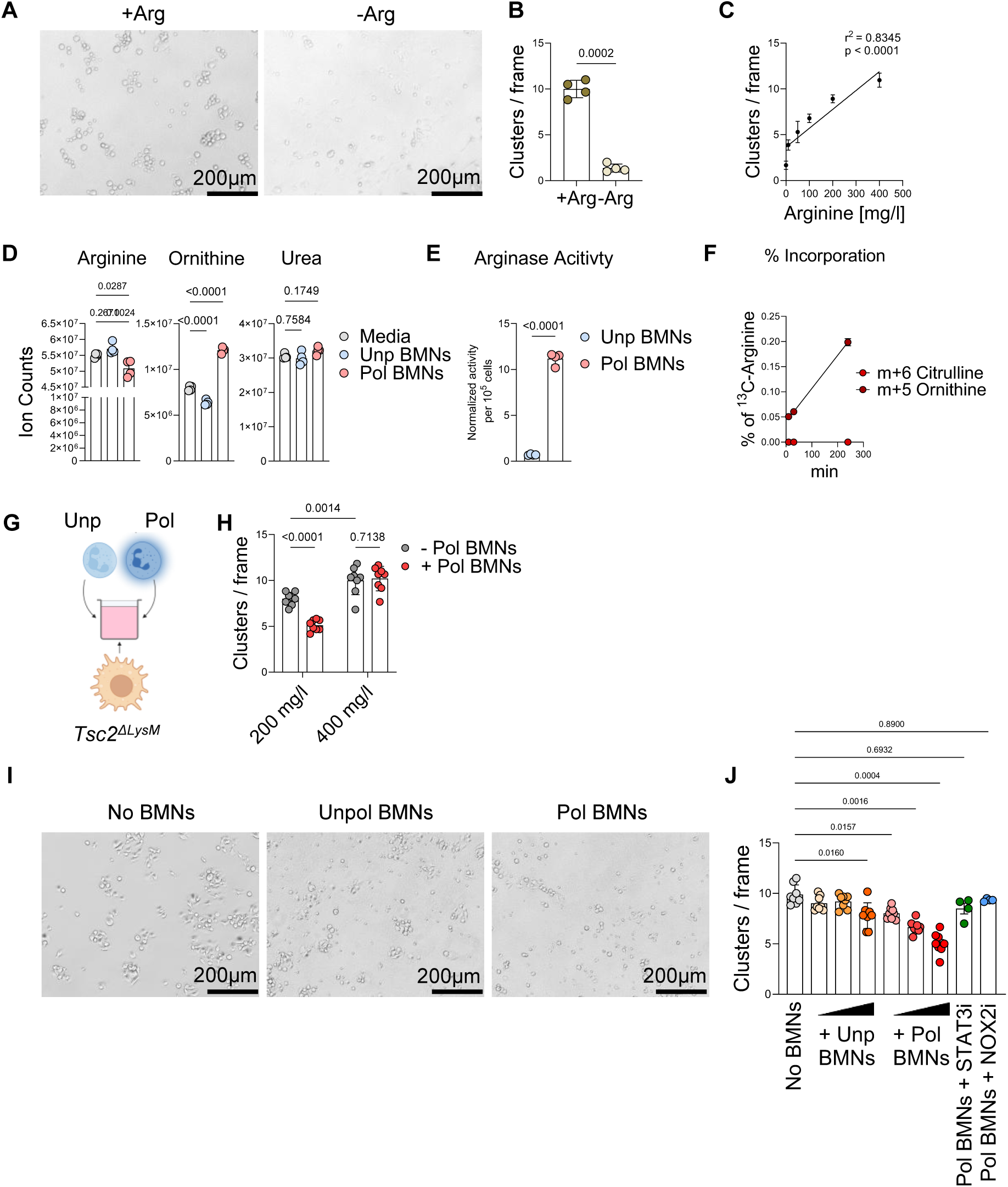
Arginine availability controls neutrophil-mediated restriction of macrophage clustering. (A, **B)** Arginine dependency of macrophage clustering. (**A**) Representative images and **(B)** quantification of cluster formation by *TSC2^ΔLyz2^* bone marrow-derived macrophages cultured in the presence (+Arg) or absence (-Arg) of arginine (n = 4 per condition). **(C)** Dose-dependent clustering by *TSC2^ΔLyz2^* macrophages across increasing arginine concentrations (0 – 400 mg/l; n = 4). **(D)** Quantification of arginine, ornithine, and urea levels in culture media supernatants of Unp and Pol BMNs (n = 4). **(E)** Arginase activity in cellular extracts of Unp or Pol BMNs (n = 4). **(F)** Stable isotope tracing of m+6 citrulline and m+5 ornithine at the indicated time points after addition of ^13^C6 arginine. Data is depicted as percentage of total labeled ^13^C arginine pool (n = 3). **(G)** Experimental design of neutrophil and *TSC2^ΔLyz2^* macrophages co-cultures for cluster analysis. (**H**) Co-culture of Pol BMNs and *TSC2^ΔLyz2^* macrophages at the indication arginine concentrations (n = 8 per condition). **(I)** Representative images and **(J)** quantification of cluster formation in co-culture with Unp BMNs or Pol BMNs, with or without NOX2 and STAT3 inhibition, and at different neutrophil-to-macrophage ratios compared to a condition without neutrophils (No BMNs) (n = 4 – 8 per condition). **(B - F, H, J)** Data are shown as means ± SD. Statistical analysis: paired t test **(B, E);** linear regression **(C);** one-way ANOVA **(D);** 2-way ANOVA **(H);** mixed-effect analysis **(J)**.

Given the high expression of Arg1 in polarized neutrophils, we hypothesized that they may limit macrophage effector functions by restricting arginine. To test this, we assessed arginine metabolism in polarized BMNs. Indeed, polarized but not unpolarized BMNs markedly lowered arginine levels in the culture medium and accumulated the downstream metabolite ornithine, indicating active arginine catabolism (**Figure 4D**). Consistently, polarized BMNs exhibited strong arginase activity compared to unstimulated cells (**Figure 4E**). Stable isotope tracing of ^13^C6-arginine in polarized neutrophils further confirmed that arginine was metabolized to ornithine, whereas no substantial conversion to citrulline was detected, indicating minimal engagement of NOS-dependent arginine metabolism (**Figure 4F**). These data establish Arg1-driven arginine catabolism as a dominant pathway in polarized neutrophils.

To investigate whether neutrophils can functionally limit macrophage-driven granuloma formation through this metabolic activity, we established a co-culture system in which polarized neutrophils were cultured with *Tsc2*-deficient BMDMs. Under standard arginine concentrations (200 mg/L), polarized neutrophils significantly reduced macrophage cluster formation. This inhibitory effect was abrogated when arginine concentrations were increased (**Figure 4G; Figure S5B**), demonstrating that neutrophil-mediated suppression of macrophage clustering depends on arginine availability (**Figure 4H**).

Finally, we assessed whether the signaling pathway controlling neutrophil polarization also regulates this granuloma-limiting function. While unpolarized neutrophils showed only modest effects on macrophage clustering, polarized BMNs inhibited cluster formation in a dose-dependent manner. Importantly, inhibition of NOX2 or STAT3 significantly impaired this inhibitory effect (**Figure 4I, J**). In summary, these findings demonstrate that alternatively polarized neutrophils restrict macrophage-driven granuloma formation through an extracellular arginine-dependent mechanism.

### Neutrophil-derived Arg1 is required to restrain macrophage-driven granuloma formation

To directly test whether neutrophil-derived Arg1 mediates the granuloma-limiting effect of neutrophils *in vivo*, we generated neutrophil-specific *Arg1*-deficient mice (*Arg1^ΔNeu^*) and analyzed granuloma formation using the coverslip model (**Figure 5A**). *Arg1^ΔNe^*^u^ mice showed a strong reduction of Arg1 expression in neutrophils (**Figure 5B, C**) and displayed exacerbated granuloma formation compared to control mice (**Figure 5D, E**). Expression of other neutrophil polarization markers remained unchanged in *Arg1^ΔNeu^* mice, indicating that Arg1 expression in neutrophils is required to limit macrophage-driven inflammation *in vivo* (**Figure S6A**).

**Figure 5:**
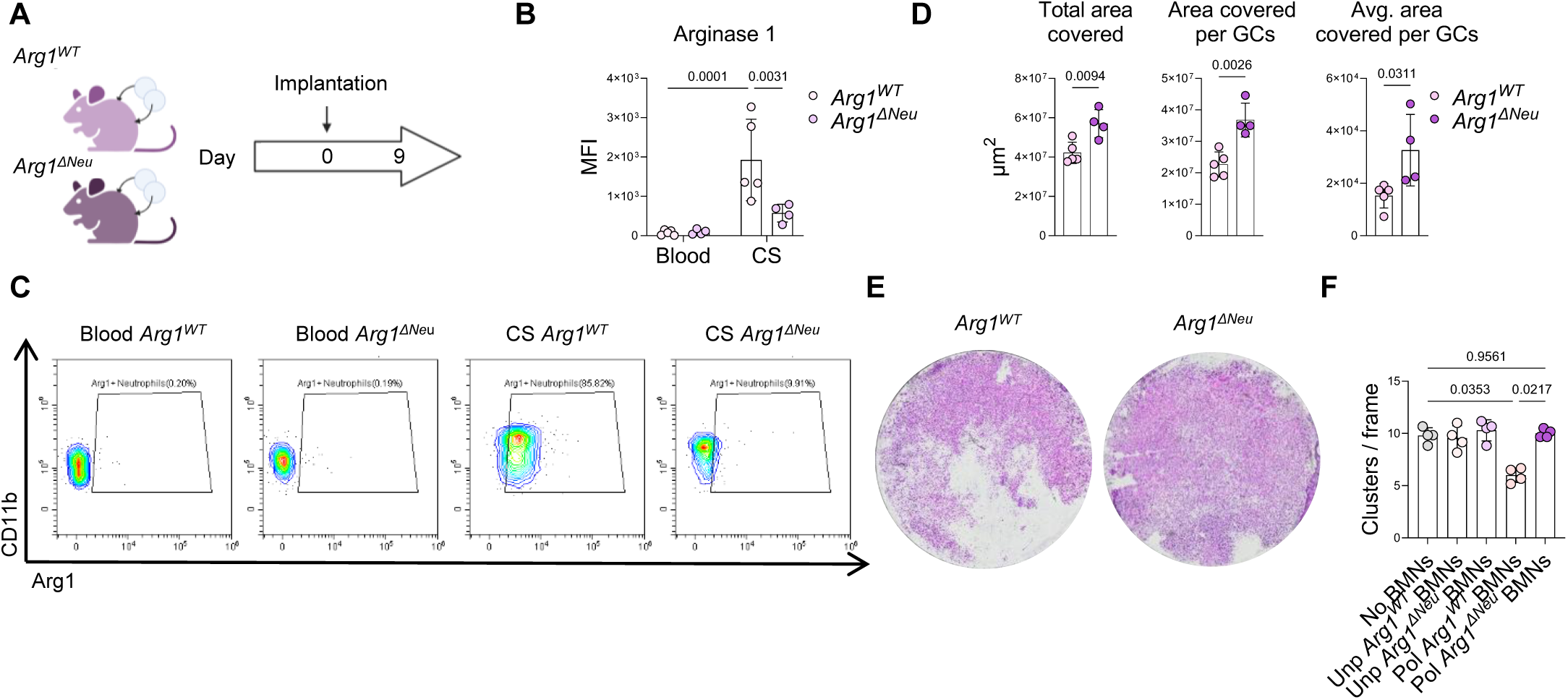
Neutrophil-derived Argq1 is required to restrain macrophage-driven granuloma formation. **(A)** Experimental design for coverslip implantation in *Arg1^WT^* and *Arg1^ΔNe^*^u^ mice. **(B)** Representative flow cytometry plots and **(C)** quantification of Arg1 in neutrophils in blood and coverslips of *Arg1^WT^* and *Arg1^ΔNeu^* mice 9 days after implantation. **(D)** Quantification and **(E)** representative images of granuloma burden in *Arg1^WT^* and *Arg1^ΔNeu^* mice at the experimental endpoint (n = 4 – 5 per genotype). **(F)** Cluster analysis of *Tsc2^ΔLyz2^* macrophages co-cultured with *Arg1^WT^* (WT) or *Arg1^ΔNeu^* (KO) neutrophils either without or with cytokine polarization (IL4 and G-CSF; n = 4 per condition). **(B, D, F)** Data are shown as means ± SD. Statistical analysis: 2-way ANOVA **(B)**; unpaired t test **(D)**; RM-one-way ANOVA **(F)**.

To further test whether neutrophil Arg1 controls macrophage granuloma formation through metabolic restriction, we again employed the *in vitro* macrophage clustering assay with *Tsc2-*deficient BMDMs. In contrast to polarized control neutrophils, polarized *Arg1^ΔNeu^* BMNs failed to suppress cluster formation (**Figure 5F, Figure S6B**). Because arginase activity generates the downstream metabolites ornithine and urea, we next tested whether these metabolites influence macrophage clustering. Addition of ornithine or urea to the culture medium did not alter cluster formation indicating that neutrophils suppress granuloma formation primarily through depletion of extracellular arginine rather than accumulation of downstream metabolites (**Figure S6C, D**). Collectively, these findings establish neutrophil-derived Arg1 as a critical regulator restraining macrophage-driven inflammation.

### Systemic arginine restriction suppresses granulomatous inflammation

We next tested whether systemic arginine restriction could therapeutically suppress granuloma formation. Two complementary approaches were used in the coverslip granuloma model: an arginine-free diet and administration of recombinant Arginase (**Figure 6A, D**). Both interventions were initiated prior to coverslip implantation as a proof-of-principle strategy. Indeed, both approaches markedly reduced granuloma formation *in vivo*, demonstrating that limiting systemic arginine availability is sufficient to suppress macrophage-driven granulomatous inflammation (**Figure 6B, C**; **Figure 6E, F**).

**Figure 6:**
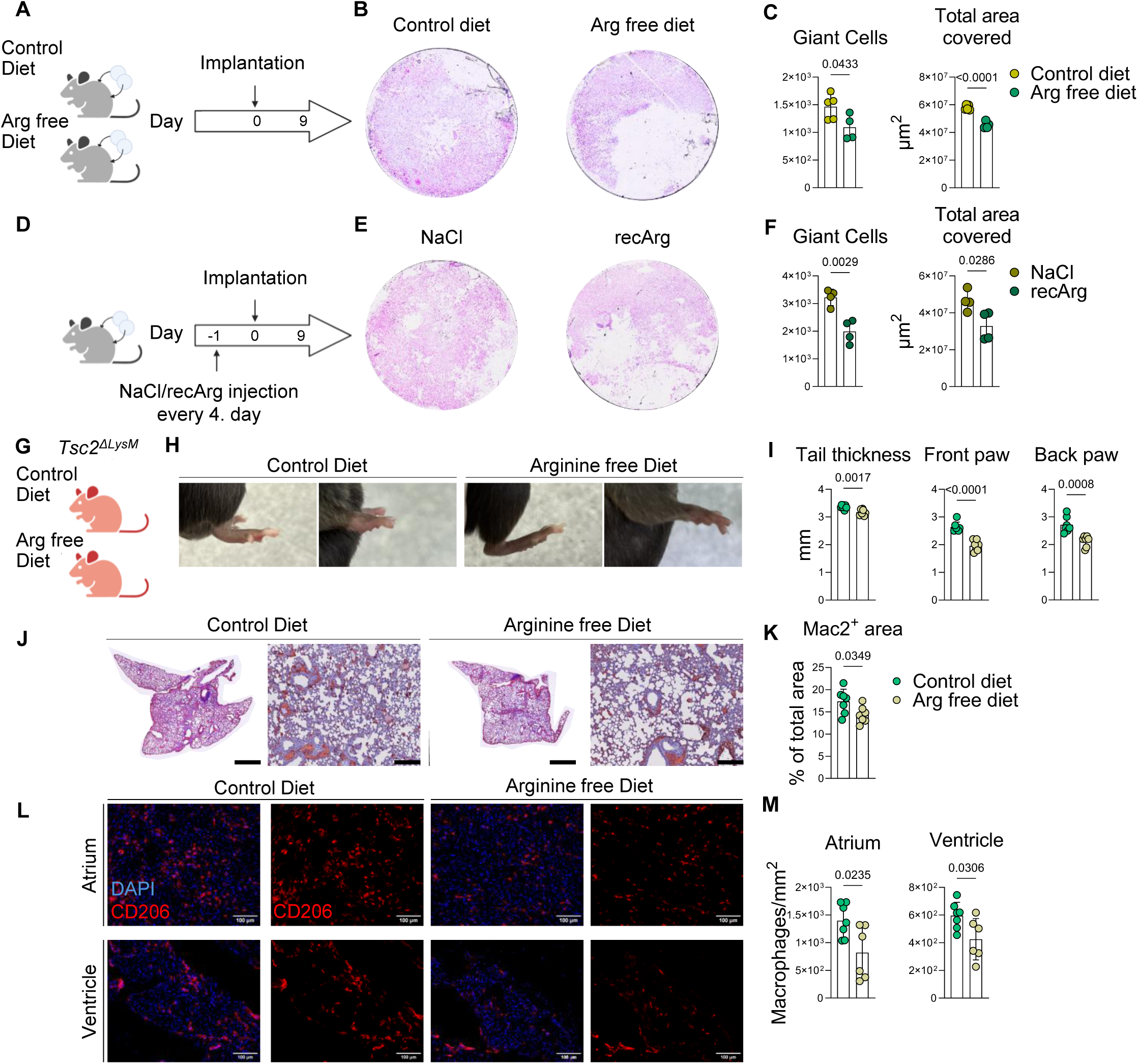
Systemic arginine restriction suppresses macrophage-driven granuloma formation *in vivo*. **(A)** Experimental design for dietary arginine restriction during coverslip-induced granuloma formation. **(B)** Representative H&E images and **(C)** quantification of granuloma burden by number of giant cells and total area covered on coverslips 9d after implantation from mice put on control or arginine free diet (n = 4 – 5 per condition). **(D)** Experimental design for enzymatic arginine depletion using recombinant arginase (recArg). **(E)** Representative H&E images and **(F)** quantification of granuloma burden by number of giant cells and total area covered on coverslips 9d after implantation from mice receiving NaCl control or recArg (n = 4 per condition). **(G)** Experimental design for dietary arginine restriction in myeloid specific *Tsc2*-deficient mice (*Tsc2^ΔLyz2^*). **(H)** Representative images of front and back paws of *Tsc2^ΔLyz2^* mice on control or arginine free diet. **(I)** Assessment of tail, front, and back paw thickness of *Tsc2^ΔLyz2^* mice on control or arginine free diet (n = 6 – 7 per condition). **(J)** Representative IHC images and **(K)** quantification of granuloma burden assessment by Mac2+ area in *Tsc2^ΔLyz2^* on control or arginine free diet (n = 6 – 7 per condition). Scale bar in the overview: 2 mm. Scale bar in the close up: 200 μm. **(L)** Representative images and **(M)** quantification of granuloma burden by CD206+ macrophages in the cardiac atrium and ventricle in *Tsc2^ΔLyz2^* on control or arginine free diet (n = 6 per condition). **(C, F, I, K, M)** Data are shown as means ± SD. Statistical analysis: unpaired t test **(C, F, I, K, M).**

To evaluate whether this metabolic intervention is effective in a more disease-relevant setting, we next examined mice with a myeloid-specific deficiency of *Tsc2*, which develop granulomatous inflammation in the lungs, heart, and paws *in vivo* that closely resembles human sarcoidosis (**Figure 6G**)^44–46^. Mice were placed on an arginine-free diet after disease onset and maintained on the diet for five weeks. Arginine restriction significantly reduced granulomatous burden in the paws (**Figure 6H, I**) and reduced granulomatous inflammation in the lungs and hearts (**Figure 6J-M**). Together, these findings demonstrate that systemic arginine restriction can suppress macrophage-driven granulomatous inflammation *in vivo*, supporting the translational potential of targeting arginine metabolism.

### Human neutrophils exhibit conserved ARG1-dependent regulatory functions

Finally, we investigated whether the regulatory features of neutrophils in mice are conserved in humans. Interestingly, neutrophils isolated from healthy human donors showed high basal expression of ARG1 in peripheral blood, indicating that arginine metabolism may represent an intrinsic feature of human neutrophils (**Figure 7A, B**). We next assessed whether arginase activity in human neutrophils is regulated through signaling pathways like those identified in mice. Human neutrophils were stimulated with phorbol 12-myristate 13-acetate (PMA) and arginase activity was measured in culture supernatants. Pharmacological inhibition of STAT3 or NOX2 significantly reduced arginase activity indicating a conserved regulatory signaling pathway (**Figure 7C**). Functionally, neutrophils from healthy donors strongly suppressed clustering of *Tsc2*-deficient macrophages in the *in vitro* clustering assay, demonstrating that human neutrophils share the capacity to limit macrophage-driven granuloma formation (**Figure 7D–F**).

**Figure 7:**
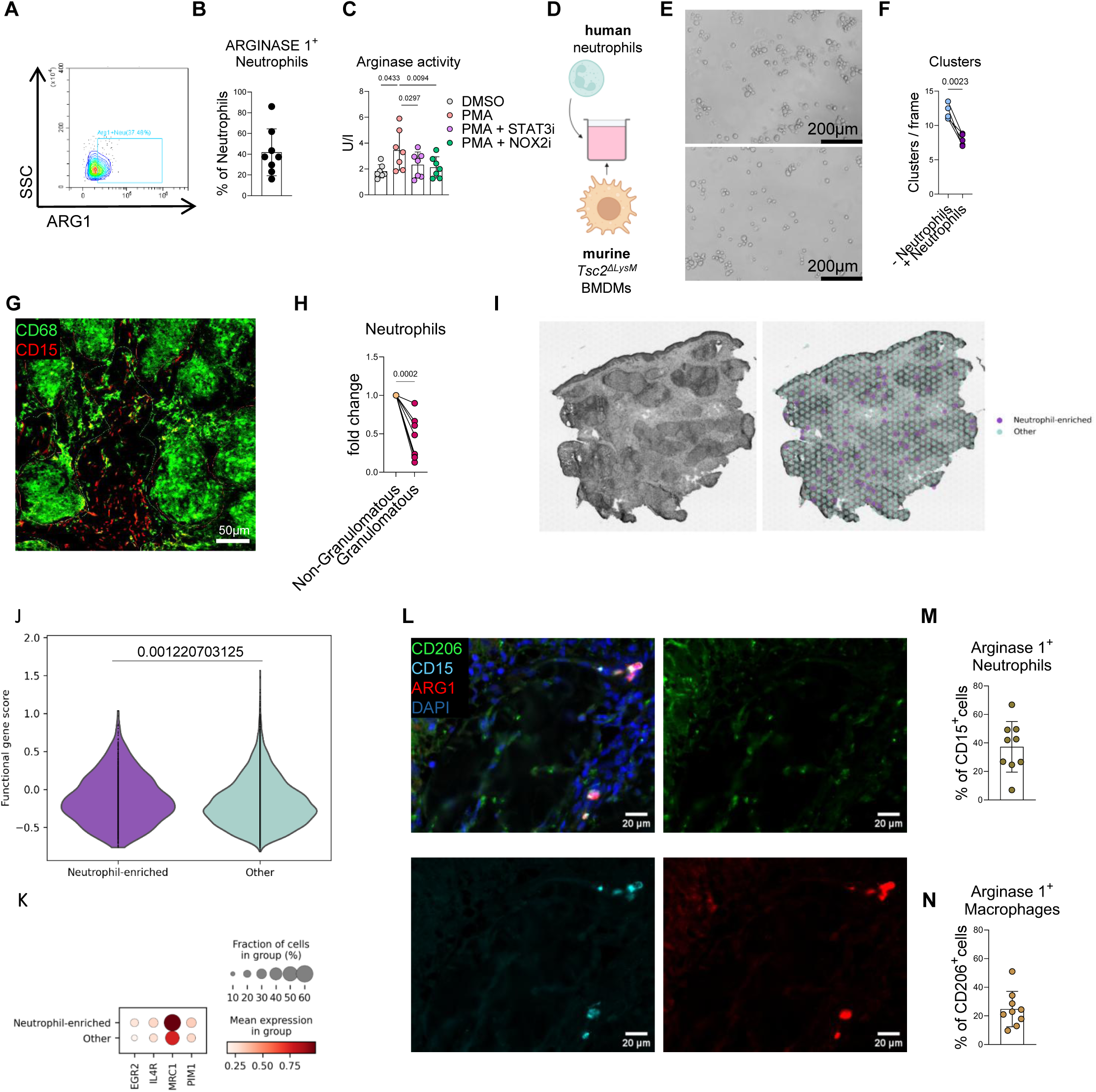
Human neutrophils exhibit a conserved alternative phenotype limiting macrophage-driven inflammation. **(A)** Representative flow cytometry plot and **(B)** quantification of ARGINASE 1^+^ neutrophils from the blood of healthy donors (n = 7). **(C)** Arginase activity in the supernatant of neutrophils stimulation with PMA in combination with STAT3 or NOX2 inhibitors (n = 6). **(D)** Schematic of co-culture system using human neutrophil and murine *Tsc2^ΔLyz2^* macrophages. **(E)** Representative images and **(F)** quantification of macrophage clustering (n = 5). **(G)** Representative immunofluorescence image and quantification **(H)** of neutrophil localization (CD15+ cells) in the non-granulomatous space or granulomatous region (CD68+ cells; n = 7). **(I)** H&E staining of a biopsy of sarcoidosis patient and spatial expression of neutrophil-enrichment. **(J)** Quantification of functional genes in neutrophil-enriched and other regions. **(J)** Dot plot depiction of functional genes of alternative polarization in neutrophil-enriched and other region. (**L**) Representative immunofluorescence image and quantification of ARG1+ neutrophils (CD15+, ARG1+; **M**) and ARG1+ macrophages (CD206+, ARG1+; **N**) in cutaneous sarcoidosis samples (n = 10). (**B, C, M, N**) Data are shown as means ± SD. Statistical analysis: RM one-way ANOVA (**C**), paired t-test (**F, H**), Wilcoxon test with Benjamini-Hochberg correction (**J**)

We next asked whether this regulatory phenotype is present in human disease. To assess the spatial relationship between neutrophils and granulomatous inflammation, we analyzed skin biopsies from patients with sarcoidosis. Neutrophils were primarily localized in skin regions lacking granulomas, consistent with the possibility that neutrophils spatially restrict granuloma formation (**Figure 7G, H**). To assess alternative polarization in this context, we analyzed our previously published spatial RNA sequencing data from skin lesions of sarcoidosis patients (**Figure 7I**)^47^. Neutrophil-associated transcripts and alternative polarization markers were enriched in tissue areas surrounding granulomatous structures rather than within the granulomas themselves (**Figure 7J, K**). Moreover, neutrophils from sarcoidosis patients exhibited marked ARG1 expression, comparable to neutrophils in the murine granuloma model (**Figure 7L - N**). Together, these data support the concept that neutrophils may spatially restrict macrophage-driven granulomatous inflammation in human tissue.

Finally, we assessed publicly available transcriptomic datasets to further support the conserved polarization program in macrophage-driven disease^48^. Analysis of human polarization markers^24^ in patients with MAS revealed increased expression of multiple alternative polarization markers (**Figure S7A**). Given the regulatory role of NOX2 in controlling ARG1 expression, we also analyzed publicly available RNA sequencing data from neutrophils of patients suffering from chronic granulomatous disease (CGD)^49^. Treatment with interferon-gamma, which is used to restore neutrophil effector functions in CGD patients, was associated with increased ARG1 expression (**Figure S7B**)^50^. Overall, our findings identify Arg1-dependent arginine depletion as a mechanism through which neutrophils restrain macrophage-driven inflammation and suggest that targeting arginine metabolism may provide a therapeutic strategy for these diseases.

## Discussion

Chronic inflammatory circuits are commonly conceptualized as self-amplifying networks in which innate immune cells perpetuate tissue damage through sustained activation. Here, we identify a mechanism through which neutrophils restrain macrophage-driven inflammation via Arg1-dependent arginine depletion. While neutrophils are increasingly appreciated as regulators of tissue homeostasis beyond their classical role in pathogen clearance^14,51,52^, their direct role in limiting macrophage-driven pathology during chronic inflammation has remained unclear.

Across complementary models of granuloma formation and MAS, we show that neutrophils accumulate early and persist throughout disease progression, where they function to constrain macrophage expansion and tissue damage. Both depletion and genetic impairment of neutrophil effector pathways exacerbated disease, whereas increased neutrophil abundance was protective. Together, these findings establish neutrophils as active regulators of macrophage-driven inflammatory responses during chronic inflammation.

Mechanistically, tissue-infiltrating neutrophils acquired a polarization state closely resembling recently described alternatively activated, pro-regenerative neutrophils^23,24^. This state is characterized by high Arg1 expression, altered maturation markers, and partial proliferative capacity, and is conserved across inflammatory contexts. While alternatively polarized neutrophils were previously linked to neuronal regeneration, the signaling pathways controlling acquisition of this phenotype remained poorly defined. Here, we identify an IL4Rα–STAT3–NOX2 signaling axis that controls this program. While STAT3 has predominantly been linked to maturation^53^ and pro-tumor functions^42^ of neutrophils, our data indicate that STAT3 also regulates Arg1-dependent neutrophil functions in inflammatory disease. Downstream of STAT3, NOX2-dependent ROS production promotes Arg1 expression, linking classical neutrophil effector pathways with the metabolic phenotype observed in polarized neutrophils^54^. Importantly, neutrophil-specific deletion of STAT3 exacerbated granulomatous inflammation *in vivo*, demonstrating functional importance of this pathway in restraining macrophage-driven pathology.

Arginine metabolism has emerged as a central regulator of immune cell function, particularly in macrophages and T cells, where it controls proliferation, polarization, and effector responses^55^. In macrophages, Arg1 has primarily been studied as a cell-intrinsic regulator that limits nitric oxide production and tissue damage^56,57^. In contrast, our data identify a mechanism in which neutrophil-derived Arg1 depletes extracellular arginine to suppress macrophage clustering and granuloma formation. This is supported by neutrophil-specific genetic ablation of Arg1, direct measurements of arginine consumption, isotope tracing, and functional rescue by arginine supplementation. Notably, downstream metabolites such as ornithine or urea did not recapitulate this effect, indicating that nutrient depletion rather than metabolite accumulation is the dominant regulatory mechanism.

Arginine-dependent metabolic regulation has been extensively studied in tumor microenvironments^58,59^, where nutrient availability influences interactions between cancer and immune cells including neutrophils^55,60–62^. Whether similar metabolic constraints regulate interactions between innate immune cell populations during chronic inflammation has remained largely unexplored. Our data provide direct genetic and functional evidence that neutrophil-derived Arg1 limits macrophage-driven inflammation through extracellular arginine depletion *in vivo*. Although Arg1 expression in neutrophils has been described, *in vivo* genetic evidence for a neutrophil-intrinsic role of Arg1 in regulating inflammatory responses has been limited^58^. Our study establishes neutrophil-derived Arg1 as a key regulator of macrophage-driven inflammatory responses.

The relevance of this regulatory circuit is supported by human data. Human neutrophils exhibit high basal ARG1 activity, conserved regulation through STAT3 and NOX2, and a potent capacity to suppress macrophage clustering *in vitro*. Given that neutrophils represent the most abundant circulating immune cell population in humans, these findings suggest that neutrophil-mediated arginine metabolism may substantially contribute to the regulation of macrophage-driven inflammation in human tissues. Spatial analysis of sarcoidosis tissue revealed that neutrophils localize preferentially to interstitial, non-granulomatous regions surrounding granulomatous lesions, and spatial transcriptomics demonstrated enrichment of neutrophil-associated transcripts and polarization markers in non-granulomatous areas. These observations are consistent with a model in which neutrophils spatially constrain macrophage expansion rather than participating within granulomas themselves. In addition, analysis of neutrophils from patients with chronic granulomatous disease (CGD) revealed that interferon-γ treatment, which restores neutrophil effector functions, is associated with increased ARG1 expression, suggesting that this pathway may be therapeutically inducible.

Moreover, the regulation of ARG1 by NOX2, together with exacerbated granuloma formation in *Ncf1*-dysfunctional mice, provides an alternative interpretation of granuloma formation in CGD. Beyond impaired microbial clearance, NOX2 deficiency may disrupt a neutrophil-intrinsic regulatory circuit that limits macrophage-driven inflammation via Arg1-dependent arginine depletion. Persistent inflammatory stimuli together with impaired metabolic restraint may therefore contribute to the excessive granuloma formation characteristic of CGD^30^.

Finally, our data identifies arginine availability as a therapeutically targetable regulator of macrophage-driven inflammation. Systemic arginine restriction, achieved through dietary or enzymatic approaches, was sufficient to suppress granuloma formation *in vivo*, including a sarcoidosis-like disease setting. Given the limited therapeutic options for granulomatous diseases, where a substantial fraction of patients develops a treatment-refractory stage, modulation of arginine metabolism may represent a mechanism-based strategy with potentially favorable tolerability. Whether such approaches can be combined with existing therapies, including steroids, methotrexate or mTOR inhibition^63,64^, warrants further investigation.

Collectively, our findings identify an alternatively polarized neutrophil state that restrains macrophage-driven inflammation through Arg1-dependent arginine depletion and establish a previously unrecognized metabolic mechanism regulating innate immune interactions during chronic inflammation.

## Data and code availability

Bulk RNA-sequencing data have been deposited to the NCBI GEO repository with the accession number GSE327893. Targeted metabolomics data is provided as supplemental file: Data S1. This paper does not report original code. Any additional data or information required to reanalyze the data reported in this paper are available from the lead contact upon request.

## Acknowledgements

Next-generation sequencing and an initial data analysis was performed by the Biomedical Sequencing Facility at CeMM Research Center for Molecular Medicine of the Austrian Academy of Sciences. LC-MS/MS analysis of neutrophil supernatant was performed by the Metabolomics Facility at Vienna BioCenter Core Facilities (VBCF), member of the Vienna BioCenter (VBC), Austria and funded by the Austrian Federal Ministry of Education, Science & Research and the City of Vienna. The work was supported by the following funding grants: The Austrian Science Fund (FWF) grants 10.55776/P34023, 10.55776/PAT3466825, FWF Sonderforschungsbereich F83 (10.55776/F8300) (all to TW), the Ann Theodore Foundation Breakthrough Sarcoidosis Initiative (to GS and TW), the Medical Scientific Fund of the Mayor of the City of Vienna (24148) and the FSR 2025-2027 Sarcoidosis Early Career Fellowship Grant (both to PE), the Deutsche Forschungsgemeinschaft (DFG, German Research Foundation) under Germany’s Excellence Strategy – EXC 2167/2 – 390884018, and FWF Austrian Science Fund (PAT 8019123) and LEO Foundation (LF-OC-24-001518) (both to GS). This work was also supported by the Federal Ministry of Economy, Energy, and Tourism of Austria and the National Foundation for Research, Technology, and Development of Austria to the Christian Doppler Laboratory for Chronic Inflammatory Skin Diseases. Illustrations were created using BioRender.com.

## Author contributions

The study was conceived and designed by TW and PE. Experiments were performed by PE, JLW, MM, AK, and IM. Data analysis and interpretation were performed by PE, JLW, PS, MM, and KB. Resources were provided by PE, JJ, GS, MHH, MH, and TW. The original draft of the manuscript was written by PE and TW. All authors reviewed and edited the manuscript.

## Declaration of conflict of interests

The authors declare no conflicts of interest.

## Supplemental Information

Data S1: Metabolomics data

Table S1: Demographic and clinical data on sarcoidosis patients.

## Material and Methods

### Human Samples

All samples were acquired in accordance with the Declaration of Helsinki, with written informed consent obtained from all subjects prior to examinations at the Medical University of Vienna, Austria, following approval (EK 1529/2025). Blood was collected in EDTA-coated tubes. Neutrophils were isolated from the erythrocyte containing pellet after density gradient centrifugation using Lymphoprep™ (STEMCELL™, 18061). After red blood cell lysis (NH4Cl based) purified neutrophils were used for downstream analysis.

### Animals

Wildtype C57BL/6J mice were purchased from the Core facility laboratory animal breeding and husbandry of the Medical University of Vienna or Janvier Labs. *Ncf1\*\** and wildtype littermate control mice were bred at Redoxis, Sweden. Fucci Tg mice were bred at the Core facility laboratory animal breeding and husbandry of the Medical University of Vienna. *Il4rα* KO and wildtype littermate control mice were bred at the Institute of Vascular Biology and Thrombosis Research of the Medical University of Vienna. Neutrophil-specific *Stat3* deficient mice (*Stat3^ΔNeu^*) were generated by crossing *Stat3^fl/fl^* and *Ly6g^Cre^* mice. *Stat3^ΔNeu^* and littermate *Stat3^WT^* mice (Stat3^fl/fl^ Ly6g^WT^) were bred at the animal facility of the University Hospital Essen. Neutrophil-specific Arginase 1 KO mice (*Arg1^ΔNeu^*) were generated by crossing *Arg1^fl/fl^* and *Ly6G^Cre^* mice. *Arg1^ΔNeu^* and littermate *Arg1^WT^* mice (*Arg1^fl/fl^ Ly6G^WT^*) were bred at the animal facility of the Center for Pathobiochemistry and Genetics of the Medical University of Vienna. *Tsc2^ΔLyz^*^2^ were generated by crossing *Tsc2^fl/fl^* mice with *Lyz2^Cre^* mice and were bred at the animal facility of the Core facility laboratory animal breeding and husbandry of the Medical University of Vienna. Mice were housed at a constant temperature of 21°C and controlled humidity of 45 – 65% with a light/dark cycle of 12h. They were provided with *ad libitum* food and water. Animal studies were approved by the official Austrian ethics committee for animal experiments (GZ BMBWF 2024-0.044.322, GZ BMBWF 2023-0.380.996, and GZ BMBWF 2023-0.062.018).

### Coverslip implantation model

Female wildtype C56BL/6J, *Ncf1*\*\*, *Il4rα* KO, *Stat3^ΔNeu^*, *Arg1^ΔNeu^* and their respective littermate control mice received implantations of 2-4 sterile glass coverslips of a diameter of 10 mm (Assistant, 41001110) via subcutaneous implantation of the rear flank. In total, mice received 1-2 coverslips per flank. Anesthesia was provided by intraperitoneal (*i.p.*) injections of 100 mg/kg ketamine (Ketamidor®, Richter Pharma) and 5 mg/kg xylazine (Rompun®, Bayer). Upon anesthesia mice received 5 mg/kg meloxicam (Metacam®, Boerhinger Ingelheim) via subcutaneous injections. Mice were kept on a heating operating pad (Harvard Bioscience) at 37°C throughout the whole surgical procedure under sterile conditions. Vitamin A containing eye cream (HYLO NIGHT®, URSAPHARMA) was put on the eyes for the duration of the surgical procedure for protection. The rear flanks were shaved using surgical scalpel blades (Swann-Morton®, 0224). For the coverslip implantation an incision of roughly 11 mm was made parallel to the spine using fine tip scissors. Upon removal of the subcutaneous connective tissue the coverslips were implanted, and incisions were closed using absorbable sutures (Vicryl suture 6-0, BV-1 needle, 30 cm purple, Ethicon, Cat. No. W9575) via continuous suturing technique. Vital signs were monitored throughout the surgical procedure and mice were monitored closely after surgery in accordance with FELASA guidelines. Experiments were terminated 6h, 12h, 24h, 3d, 9d, and 21d after coverslip implantation and mice were sacrificed by an *i.p.* injection of a lethal dose of ketamine and xylazine followed by cervical dislocation. In experiments requiring terminal blood withdrawal, blood was collected from anesthetized mice by cardiac puncture prior to cervical dislocation.

### Macrophage activation syndrome model

Male wildtype C56BL/6J received 5 injections of 200 μg CpG (ODN1826, InvivoGen, tlrl-1826; MCE, HY-146245) or PBS control via *i.p.* injection every second day. Mice were monitored closely after first injection in accordance with FELASA guidelines and weighed daily. Experiments were terminated 10d after the first injection and mice were terminated by an *i.p.* injection of a lethal dose of ketamine and xylazine flowed by cervical dislocation. In experiments requiring terminal blood withdrawal, blood was collected from anesthetized mice by cardiac puncture prior to cervical dislocation.

### *In vivo* neutrophil depletion

Neutrophil depletion was used in coverslip implantation model and macrophage activation syndrome model. In both experiments mice received Ly6G (InVivoMAb anti-mouse Ly6G, bioxcell, #BE0075-1) or isotype control antibody (InVivoMAb rat IgG2a isotype control, bioxcell, #BE0089) 1d before implantation or the first CpG injection respectively (400 μg per mouse) and afterwards every second day (100 μg per mouse) throughout the duration of the experiment to maintain the neutrophil depletion.

### *In vivo* G-CSF injection

Neutrophilia was induced by daily injections of 5 μg of murine recombinant G-CSF (Mouse G-CSF, Animal-Free Recombinant Protein, PeproTech®, ThermoFisher Scientific, AF-250-05). Mice received the first injection 1d before coverslip implantation and then daily injections for maintenance.

### Arginine free diet

Custom made arginine free diet and control chow diet was purchased from ssniff. For coverslip implantation experiments, mice were put on arginine free diet or the respective control diet 1 week prior to coverslip implantation and maintained on the diet for the duration of the experiment. *Tsc2^ΔLyz^*^2^ mice were maintained on arginine free or control chow diet for 5 weeks. In each experiment mice were monitored and weighed regularly.

### Recombinant arginase treatment

Recombinant arginase (BCT-100) was kindly provided by Gernot Schabbauer from the Medical University of Vienna. Mice received 50 mg/kg of recombinant arginase or respective NaCl control injections 1 day before coverslip implantation and then every 4. day for maintenance until the experimental endpoint.

### Tissue Processing

Liver and lung samples were fixed overnight using 4% formaldehyde (ROTI® Histofix, Roth P087.1) and then embedded into paraffin (Parabplast, Leica Biosystems, 39601006) using the KOS tissue processor (Milestone). Heart samples were embedded using OCT embedding matrix (Fischer Scientific, 30226281).

### May–Grünwald–Giemsa Staining

Coverslips were briefly rinsed in PBS immediately after removal and fixed in methanol for 7 min at room temperature. After air-drying (∼30 min), coverslips were mounted onto glass slides using Neo-Mount (Merck, 109106) and allowed to dry for at least 20 min. Coverslips were processed depending on implantation duration. For 9-day samples, slides were permeabilized in 3% PBS–Triton for 10 min; for 3-day samples, permeabilization was performed for 5 min; and for 1-day samples or low cell density, permeabilization was omitted. Slides were briefly rinsed in distilled water and stained with May–Grünwald solution (5–10 min; Sigma, MG500-500ml), followed by incubation in phosphate buffer (1.5 min) and staining with freshly prepared Giemsa solution (15– 25 min; Sigma GS500-500ml). Slides were briefly rinsed in distilled water (10 s). Coverslips were then gently detached, air-dried, and mounted cell-side down onto fresh slides using Neo-Mount. Images were acquired on a BX63 Intelligent Microscope (Olympus).

### Flow cytometry

Single cell suspension from coverslips was obtained using digestion with Dispase II (2.4 U/ml; Sigma D4693) in PBS + 2% FCS at 37°C for 25 min. Single cell suspension was obtained by using a 100μm strainer. For blood isolated from mice, erythrocyte lysis was performed 2x on ice (Biolegend, 420301) and cells were used for flow cytometric analysis. Single cell suspension from liver samples was obtained by using the Multi Tissue Dissociation Kit (Miltenyi, 130-110-201) as suggested by the protocol. Spleens were processed by smashing on a 70μm strainer followed by erythrocyte lysis. To block unspecific binding TruStain FcX™ PLUS (Biolegend, 156603) and Human TruStain FcX™ (Biolegend, 422301) was used. For intracellular staining Foxp3 / Transcription Factor Staining Buffer Kit (Cytek®, SKU TNB-0607-KIT) was used as suggested by the company’s protocol. For staining of phosphorylated proteins cells were fixed (ThermoFisher Scientific, 00-8222-49) for 20 min and permeabilized using ice-cold methanol for 15 min. Surface staining was performed in PBS at room temperature for 15 – 30 min. Intracellular staining was performed at 4°C overnight.

Antibodies used for the study were: Anti-Puromycin (Alexa Fluor 647; Merck, MABE343-AF647); Arginase-1 (APC; Thermo Fisher Scientific, 17-3697-82); CD101 (PC7; Thermo Fisher Scientific, 25-1011-82); CD115 (PE/Dazzle 594; BioLegend, 135527); CD11b (APC; TONBO, 20-0112); CD11b (eFluor 450; eBioscience, 48-0112-82); CD11b (FITC; BD Biosciences, 557396); CD11b (PerCP PC5.5; TONBO, 65-0112-U025); CD11b (Violet610; BioLegend, 101237); CD11c (PE-Cy7; eBioscience, 25-0114-81); CD15 (BV421; BioLegend, 323040); CD204 (PerCP PC5.5; BioLegend, 154715); CD45.2 (Alexa Fluor 488; BioLegend, 109816); CD45.2 (Pacific Blue 450; BioLegend, 109820); CD45.2 (PE; BioLegend, 10980); CD62L (APC; BioLegend, 104411); CD62L (PE; BioLegend, 104407); CXCR4 (CD184) (FITC; Invitrogen, 53-9991-80); F4/80 (APC; BioLegend, 123116); F4/80 (BV605; BioLegend, 123133); F4/80 (Violet660; BioLegend, 123149); I-A/I-E (MHC II) (BV605; BD Biosciences, 743872); iNOS (PE; eBioscience, 12-5920-80); Ly6C (APC-A700; BioLegend, 128024); Ly6C (APC-Cy7; BD Biosciences, 560596); Ly6C (PE; BioLegend, 128007); Ly6G (APC-A700; BioLegend, 127622); Ly6G (BV510; BioLegend, 127633); Ly6G (PE; BioLegend, 164503); Ly6G (PerCP-Cy5.5; BioLegend, 127615); Mouse IgG1, κ Isotype Control (PE; BioLegend, 400139); Mouse IgG2a, κ Isotype Control (PE; eBioscience, 12-4321-80); NOX2 (Alexa Fluor 488; Proteintech, CL488-19013); pSTAT3 (PE; BioLegend, 651003); Rat IgG2a, κ Isotype Control (APC; Invitrogen, 17-4321-81); Viability (7-AAD; Sigma-Aldrich, A9400); Viability (Zombie Aqua; BioLegend, 423101); Viability (fixable; eFluor 780; Thermo Fisher Scientific, 65-0865-14).

### Hematoxylin & Eosin staining

Coverslips were fixed with 4% formaldehyde (ROTI® Histofix, Roth P087.1) after explantation. After washing they were mounted (cell side up) on to glass slides using NeoMount^TM^ (Sigma-Aldrich, 109016). Staining was performed with Mayer’s hematoxylin (Merck 1051740500) and Eosin Y (Sigma-Aldrich 230251) or Eosin Y + Phloxine (Merck, HT110116-500ML). Between Hematoxylin and Eosin there was a differentiation step using 1x Scott Tap Water substitute. Coverslips were mounted onto fresh slides using NeoMount^TM^. Images were acquired on a BX63 Intelligent Microscope (Olympus) and image analysis was performed using HALO^®^ (Indica Labs). Tissue Sections (3μm) from paraffin embedded liver samples were deparaffinzed and rehydrated before Hematoxylin and Eosin staining. The staining was performed as with coverslips, with an additional differentiation step using 3% HCl (Roth, 9969.1). Images were acquired on a BX63 Intelligent Microscope (Olympus). For quantification of inflammatory foci 10 random images were taken and infiltrates counted manually^65^.

### Immunohistochemistry Staining

Sections underwent heat-induced epitope retrieval in retrieval buffer (Thermo Fisher, 00-4955-58) using autoclaving (120 °C), followed by cooling. Slides were washed in PBS and, if required, permeabilized with 0.3% Triton X-100. Endogenous peroxidase activity was quenched with 3% H₂O₂ (Roth, 9618.4). Sections were blocked with serum in PBST and incubated overnight with primary antibody (anti-Mac2; CEDARLANE, CL8942AP; 1:10 000) diluted in PBST (0.025%) containing 2.5% serum. After washing, biotin blocking was performed (Vector Labs, SP 2002), followed by incubation with biotinylated secondary antibody (Vector Labs, BA-9400) and HRP–streptavidin (Leica, RE7104-CE). Signal was developed using AEC substrate (Biolegend, 925804). Sections were counterstained with Meyer’s hematoxylin, rinsed, and mounted with Aquatex (Merck, 1085620050). Images were acquired on a BX63 Intelligent Microscope (Olympus) and image analysis was performed using ImageJ (Indica Labs).

### Bulk RNA-sequencing

Cells from blood and coverslip were isolated as described above. To obtain enough cells, two mice were pooled per sample. Neutrophils from blood and coverslips were FACS-sorted based on viable, CD11b, and Ly6G+ cells. A total of 60 000 cells were directly sorted into to Buffer RLT (Qiagen) containing beta-mercaptoethanol (1%, ThermoFisher Scientific, 21985023) and RNasin® Plus Ribonuclease Inhibitor (2 U/l, Promega, N2611). RNA was extracted using the RNeasy Micro Kit (Qiagen, 74004) according to manufacturer’s protocol.

For the preparation of NGS libraries from low-input samples we followed the Smart-seq2 protocol. The subsequent library preparation from the amplified cDNA was performed using the Nextera XT DNA library prep kit (Illumina, San Diego, CA, USA). Library concentrations were quantified with the Qubit® 1X dsDNA HS Assay Kit (Q32856, Thermo Fisher Scientific, Waltham, MA, USA) and the size distribution was assessed using the High Sensitivity DNA Kit (5067-4626, Agilent, Santa Clara, CA, USA) on a 2100 Bioanalyzer High-Resolution Automated Electrophoresis instrument (G2939A/B, Agilent). Before sequencing, sample-specific NGS libraries were diluted and pooled in equimolar amounts.

Expression profiling libraries were sequenced on a NovaSeq 6000 instrument (Illumina, San Diego, CA, USA) following a 50-base-pair, paired-end recipe. Raw data acquisition (NovaSeq Control Software, NCS, 1.8.1) and base calling (Real-Time Analysis Software, RTA, 3.4.4) was performed on instrument, while the subsequent raw data processing off the instruments involved two custom programs based on Picard tools (2.19.2). In a first step, base calls were converted into lane-specific, multiplexed, unaligned BAM files suitable for long-term archival (IlluminaBasecallsToMultiplexSam, 2.19.2-CeMM), thereby annotating an eventual start of NGS adapter sequences in XT tags. In a second step, archive BAM files were demultiplexed into sample-specific, unaligned BAM files (IlluminaSamDemux, 2.19.2-CeMM), annotating unique molecular index (UMI) sequences in RX tags and UMI base quality scores in QX tags.

NGS reads were mapped to the Genome Reference Consortium GRCm39 assembly via “Spliced Transcripts Alignment to a Reference” (STAR, 2.7.11b) utilising the “basic” GENCODE transcript annotation from version vM35 (May 2024) as reference transcriptome. Since the mm39 assembly flavour of the UCSC Genome Browser was preferred for downstream data processing with Bioconductor packages for entirely technical reasons, GENCODE transcript annotation had to be adjusted to UCSC Genome Browser sequence region names. STAR was run with options recommended by the ENCODE project. Metadata annotation, such as the start of NGS adapter sequences (in XT tags), were propagated from unaligned BAM files to the aligned BAM files via Picard MergeBamAlignment and duplicated alignments were marked with Picard MarkDuplicates in a UMI-aware manner but not removed. The differential expression modelling was based on R (4.4.2) and Bioconductor (3.20). NGS read alignments overlapping Ensembl exon features were counted with the Bioconductor GenomicAlignments (1.42.0) package via the summarizeOverlaps function in Union mode, ignoring secondary alignments, alignments not passing vendor quality filtering and for UMI-supporting protocols only, also duplicate alignments. Since the Smart-seq2 protocol is not strand specific, all alignments irrespective of the feature (i.e., gene, transcript, and exon) orientation were counted. Exon-level counts were aggregated to gene-level counts and the Bioconductor DESeq2 (1.46.0) package was used to test for differential expression based on a model using the negative binomial distribution.

An initial exploratory analysis included principal component analysis (PCA), multi-dimensional scaling (MDS) and sample distance heatmap plots (ggplot2, 3.5.1), all based on a variance-stabilising transformation (DESeq2) and annotated with variables used in the expression modelling.

Biologically meaningful contrasts were extracted from the model, log2-fold values were shrunk with the CRAN ashr (2.2.-63) package, while two-tailed p-values obtained from Wald testing were adjusted with the Bioconductor Independent Hypothesis Weighting (IHW, 1.34.0) package. The contrasts were visualised via MA, volcano (Bioconductor EnhancedVolcano, 1.24.0) and expression heatmap (Bioconductor ComplexHeatmap, 2.22.0) plots based on hierarchical clustering method “complete” of Euclidean distances. The resulting gene tables were annotated and subsequently filtered for significantly differentially up- and down-regulated genes and independently subjected to gene set enrichment analysis (Enrichr).

### Differential gene expression analysis, PCA plot, heatmap and volcano plot generation

Genes with lower than 10 counts across the samples were not included in the analysis. DEseq2 (v1.42.1) was used to normalize the raw counts and to perform differential gene expression analysis. Variance stabilizing transformation (vst) function, from DEseq2 package, was applied to the raw counts to transform the data for Principal Component Analysis. Based on the variable ‘Condition’, plotPCA function was applied to the transformed data to visualize the global transcriptomic difference among the samples. Biomart (v2.58.2) was used to convert the Ensemble ID to Gene ID, the significant genes were filtered using the threshold Adjusted P value <0.05 and Log2Foldchange >0.5. Genes of interests among the significant genes were extracted from the DEseq2 object using the ‘count’ function. Z-scores were obtained from Expression values after Log2 transformation and standardization per gene across samples. The heatmaps were generated using ComplexHeatmap package (v2.18.0) in R(v4.3.0) from the z-scores. Volcano plots were generated using ggplot2 (v3.5.1).

### Gene Set Enrichment Analysis (GSEA) and Gene Ontology (GO) analysis

The hallmark gene sets along with other genes sets of interests were obtained from msigdbr package (v10.0.1) for Mus musculus. Following that GSEA was performed on the pre-ranked genes using a function, ‘GSEA’, from clusterprofiler package(v4.10.1). Both GSEA and GO enrichment plots were generated using ggplot2.

### SCENITH^TM^

The protocol was adapted from previous work^66^. Single cell suspension from blood and coverslip was obtained as for flow cytometry described above. 1x10^5 cells per condition were plated in a 96 well plate and incubated for 10 min at 37°C. Afterwards cells were incubated with 2-Deoxy-D-glucose (2DG, Merck, D8375-10MG), oligomycin A (MCE, MEXHY-16589-1), a combination of 2DG and oligomycin, or DMSO control for 10 min followed by incubation with puromycin (Merck, P7255) for 30 min at 37°C. Reaction was stopped by adding 100 µl of ice cold PBS and cells were stained with desired surface antibodies, fixed with Foxp3 / Transcription Factor Staining Buffer Kit (Cytek®, SKU TNB-0607-KIT) and stained overnight with anti-puromycin antibody (Merck, MABE343-AF647).

### Murine neutrophil isolation and *in vitro* polarization

Neutrophils were isolated from the bone marrow of mice by flushing with PBS and subsequent erythrocyte lysis (Biolegend, 420301). To purify neutrophils the EasySep™ Mouse Neutrophil Enrichment Kit (Stemcell™, 19762) was used according to manufacturer’s protocol. To polarize neutrophils, cells were seeded at a density of 1x10^6^/ml in RPMI 1640 medium (ThermoFisher Scientific, 21875-034) supplemented with fetal bovine serum (FBS, ThermoFisher Scientific, A5669801) and Penicillin-Streptomycin (ThermoFisher Scientific, 15140122). For polarization cells were treated with murine recombinant IL-4 (25 ng/ml; ThermoFisher Scientific, 214-14) and G-CSF (25 ng/ml; ThermoFisher Scientific, 250-05) for up to 24 hours depending on the experimental set-up. For pharmacological inhibitor studies following compounds were used: WP1066 (STAT3i; 5 – 10 µM; MCE, HY-15312), GSK2795039 (NOX2i; 5 – 20 µM; MCE; HY-18950), and N-acetyl-cysteine (NAC; 500 µM, Merck, A9165-5G).

### Bone marrow-derived macrophage generation and *in vitro* clustering

To generate bone marrow-derived macrophages from *Tsc2^ΔLyz2^* mice, bone marrow was flushed using PBS followed by erythrocyte lysis (Biolegend, 420301). Cells were plated on 10 cm non-TC treated dishes in RPMI 1640 medium (ThermoFisher Scientific, 52400-025) supplemented with fetal bovine serum (FBS, ThermoFisher Scientific, A5669801), Penicillin-Streptomycin (ThermoFisher Scientific, 15140122), and murine recombinant M-CSF (30 ng/ml; Biolegend, 576406) for 7 days. On day 3 after seeding fresh M-CSF was added. After differentiation, BMDMs were detached using Enzyme Free Cell Dissociation Solution PBS Based (Merck, S-014-B) and re-seeded for cluster analysis at a density of 1 x 10^5^ cells/ml in the same medium as for differentiation supplemented with 20 ng/ml M-CSF for 4 – 5 days. For cluster analysis in arginine-free media RPMI 1640 Medium for SILAC (ThermoFisher Scientific, 88365) supplemented with Penicillin-Streptomycin (ThermoFisher Scientific, 15140122), dialyzed Fetal Bovine Serum (ThermoFisher Scientific, 26400044), L-Lysine-monohydrochloride (Merck, L8662-100G), and L-Arginine (ranging from 0 – 400 mg/l; Merck, A8094-25G). For co-culture experiments human neutrophils were added at a ratio of 1:1 3 days after re-seeding of the BMDMs. In the case of co-cultures with murine neutrophils, they were either polarized (in the presence or absence of STAT3 and NOX2 inhibitors) or unpolarized for 24h and added to the BMDMs 3-4 days after re-seeding. Treatment with L-Ornithine monohydrochloride (500 µM; Merck, O6503-25G) or Urea (500 µM; Merck, U5378-100G) was started directly after reseeding. Clusters were imaged at 6 random spots per well using an Olympus IC51 inverted microscope and quantified as average clusters per field of view.

### LC-MS based metabolomics

Control medium, and supernatant from unpolarized or polarized neutrophils after 24h of incubation were collected, centrifuged once (500g, 5 min) and snap-frozen using liquid nitrogen. The samples were analyzed by liquid chromatography directly coupled to tandem mass spectrometry (LC-MS/MS), using both reversed phase chromatography (RP) and hydrophilic interaction liquid chromatography (HILIC). Before RP-LC-MS/MS analysis, 100 µl of the extracts were dried down in a vacuum centrifuge and were resolved in 100 µl 0.1% formic acid in water. Detection and quantification were done by LC-MS/MS using an Ultimate 3000 HPLC system (Dionex, Thermo Fisher Scientific), directly coupled via electrospray ionization to a TSQ Altis mass spectrometer (Thermo Fisher Scientific). 1 µl of each resolved sample was injected onto a Kinetex (Phenomenex) C18 column (100 Å, 150 x 2.1 mm) connected with the respective guard column, employing a 6-minute-long linear gradient from 100% A (1 % acetonitrile, 0.1 % formic acid in water) to 90% B (0.1 % formic acid in acetonitrile) at a flow rate of 80 µl/min. Employing selected reaction monitoring (SRM), the following transitions were applied in the positive ion mode: m/z 61 to m/z 44 (urea), m/z 76 to m/z 30 (glycine), m/z 89 to m/z 72 (putrescine), m/z 116 to m/z 70 (proline), m/z 132 to m/z 90 (creatine), m/z 133 to m/z 70 (ornithine), m/z 134 to m/z 74 (aspartate), m/z 146 to m/z 72 (spermidine), m/z 150 to m/z 133 (methionine), m/z 175 to m/z 70 (arginine), m/z 176 to m/z 159 (citrulline), m/z 205 to m/z 188 (tryptophan), m/z 203 to m/z 112 (spermine), m/z 289 to m/z 271 (arginosuccinate) and 399 to m/z 250 (SAM). In an independent experiment using HILIC, 1 µl of the extracted sample was directly injected onto a polymeric iHILIC-(P) Classic HPLC column (HILICON, 100 x 2.1 mm; 5 µm) and the respective guard column, operated at a flow rate of 100 µl/min. A linear gradient (A: acetonitrile; B: 10 mM aqueous ammonium bicarbonate, supplemented with 0.1 µg/ml medronic acid) starting with 25% B and ramping up to 75% B in 9 minutes was used for separation. Using a TSQ Quantiva mass spectrometer (Thermo Fisher Scientific), the following SRM transitions were used for quantitation in the negative ion mode: m/z 124 to m/z 80 (taurine), m/z 133 to m/z 115 (malate), m/z 165 to m/z 147 (phenyllactic acid) and m/z 175 to m/z 85 (aconitate). Additionally, SRM transitions were used for quantitation in the positive ion mode: m/z 118 to m/z 101 (guanidinoacetate), m/z 118 to m/z 72 (valine), m/z 131 to m/z 72 (agmatine), m/z 132 to m/z 86 (isoleucine & leucine), m/z 147 to m/z 30 (glutamine), m/z 148 to m/z 84 (glutamic acid), m/z 118 to m/z 101 (guanidinoacetate), m/z 205 to m/z 188 (tryptophan) and m/z 385 to m/z 136 (SAH). In all cases, authentic standards have been used for determining optimal collision energies of the SRM transitions and for validating experimental retention times via standard addition to a pooled quality control sample. The data interpretation was performed using TraceFinder (Thermo Fisher Scientific).

### Arginase activity assay

For mouse neutrophils, cells were polarized for 24h or left unpolarized *in vitro* as outline above. After 1 wash with PBS, the cellular protein fraction was isolated using CelLytic™ MT Cell Lysis Reagent (Merck, C3228-50ML) according to manufacturer’s protocol. The cellular protein faction was then used for an Arginase Activity Assay Kit (Merck, MAK112-1KT), which was performed according to manufacturer’s protocol.

For human neutrophils, cells were isolated as described above and seeded on TC-treated 12-well plates at a density of 2 x 10^5^/ml in RPMI 1640 Medium for SILAC (ThermoFisher Scientific, 88365) supplemented with Penicillin-Streptomycin (ThermoFisher Scientific, 15140122), dialyzed Fetal Bovine Serum (ThermoFisher Scientific, 26400044), and L-Lysine-monohydrochloride (Merck, L8662-100G). After 30 min of time to settle, cells were treated with WP1066 (10 µM), GSK2795039 (20 µM), or corresponding DMSO control for 30 min before stimulation with Phorbol 12-myristate 13-acetate (PMA; Merck, 524400-1MG) for 2h. Supernatant was isolated and centrifuged once followed by snap-freezing in liquid nitrogen. A fraction of the supernatant was subsequently used for the arginase activity assay according to manufacturer’s protocol.

### Stable isotope tracing and metabolite extraction

Cells were incubated for 16h as described above with IL-4 and G-CSF. Media was removed and replaced with arginine-free media as outlined above with 1.15 mM ^13^C6-labeled arginine tracer (Sigma, 643440) for defined time periods (10 min, 30 min, and 4 h).

For metabolite extraction, 20 µL of sample (cell suspension or medium solution) was added to 80 µL of ice-cold methanol containing 0.5% (v/v) formic acid. Samples were incubated on dry ice for 3 min to ensure rapid quenching of metabolic activity. Subsequently, extracts were neutralized by adding 8 µL of 15% (w/v) aqueous ammonium bicarbonate solution and incubated on dry ice for an additional 20 min. Samples were then centrifuged at 14,000 × g for 20 min at 4 °C. The supernatants were transferred to LC-MS vials for analysis.

Metabolite analysis was performed using a Vanquish Flex UHPLC system (Thermo Fisher Scientific) coupled to an Orbitrap Exploris 240 mass spectrometer (Thermo Fisher Scientific). Chromatographic separation was achieved using a XBridge BEH Amide column (2.1×150 mm, 2.5 µm; Waters). Solvent A consisted of 95:5 (v/v) water:acetonitrile containing 20 mM ammonium acetate, 20 mM ammonium hydroxide, and 5 µM ammonium hydrogen phosphate, pH 9.5, and Solvent B was acetonitrile. The flow rate was set to 0.15 mL/min, the column temperature was kept at 25°C and the autosampler was maintained at 4°C. The gradient elution program was as follows^67^: 0– 2 min, 90% B; 2–3 min, 90% to 75% B; 3–7 min, 75% B; 7–8 min, 75% to 70% B; 8–9 min, 70% B; 9–10 min, 70% to 50% B; 10–12 min, 50% B; 12–13 min, 50% to 25% B; 13–14 min, 25% B; 14–16 min, 25% to 0% B; 16–20.5 min, 0% B; 20.5–21 min, 0% to 90% B; 21–25 min, 90% B. Injection volumes were 10 µL for cell extracts and 2 µL for media samples.

Mass spectrometric detection was performed in positive electrospray ionization mode. The spray voltage was set to 3.2 kV. The ion transfer tube temperature was maintained at 300 °C, and the vaporizer temperature was set to 30 °C. Sheath gas, auxiliary gas, and sweep gas were set to 35, 10, and 0.5 arbitrary units, respectively. The RF lens was set to 70%. Full scan MS data were acquired at a resolution of 180,000 over a mass range of m/z 70–1000. The automatic gain control (AGC) target was set to 1000%, with a maximum injection time of 500 ms.

Raw LC-MS data files (.raw) were converted to mzXML format using MSConvertGUI (ProteoWizard). Data were processed using El-MAVEN (v0.12.0) for metabolite identification and isotopologue analysis.

### Immunocytochemistry

5µm acetone-fixed cryosections of skin punch biopsies from patients affected by steroid-refractory cutaneous sarcoidosis were de-frosted for 10 min in 1x PBS and then blocked for 20 min with 2% goat serum in 2% BSA/PBS. Slides were stained with a mixture of directly conjugated antibodies specific to CD68-APC (Biolegend, 333810) and CD15-PE (Biolegend, 301906) in 2% BSA/PBS for 1h at room temperature and subsequently washed 3 x 5 min in 1x PBS. After washing, the slides were counterstained with 4,6-Diamidino-2-phenylindole (DAPI; Roche Applied Science, 10236276001), washed 1 x 5 min with PBS and mounted in Fluoromount-G™ Mounting Medium (ThermoFisher Scientific). Samples were scanned using an Z1 Axio Observer microscope equipped with an LD Plan-Neofluar 20x/0.4 objective (Zeiss) connected to a TissueFAXS imaging system and analyzed using TissueQuest software (TissueGnostics). Isotype-matched negative controls were done in parallel using directly conjugated antibodies with the same concentration.

Alternatively, 5 µm thick paraffin sections were cut on a microtome and dried overnight, or 5 µm thick OCT-embedded cryosections were cut from a cryostat. Deparaffinization was performed in Neo-Clear™ (Merck, 1.09843) an aliphatic hydrocarbon mixture solution, and a series of ethanol washes (96% and 100%) and rehydrated in dH2O and 1x PBS. Heat-induced epitope retrieval was performed at pH 6 or pH 8 antigen retrieval solution (Thermo Scientific, 00-4955-58, 00-4956-58) in boiling solution for 30 minutes. Frozen sections were brought to room temperature and briefly washed in PBS. Both types of sections were then blocked in serum and incubated overnight with a primary antibody cocktail. The following day, sections were washed in 1x PBS containing 0.025% Triton-X (Sigma, X100) and incubated with a fluorescently labelled secondary antibody mixed with DAPI nuclear stain solution (Merck, D9542) for 1 hour at room temperature. Slides were then washed, mounted, and imaged. Antibodies used were Anti-CD206 1:100 (Thermo Fisher, PA5-46994), Anti-CD15 1:100 (Thermo, MA1-022), Anti-ARG1 1:200 (CST, 93668), and Anti-Mouse AF555, (Thermo Fisher, A31570), anti-Rabbit, (Thermo Fisher, A32795), and Anti-Goat IgG H&L Alexa Fluor® 488, (Thermo Fisher, A32814; all 1:500).

Immunofluorescence images were acquired on an inverted light microscope APEX, model APX100 (Evident, Germany). Whole-well overviews were taken at 200x magnification. Image analysis was performed in QuPath v0.5. In short, well areas containing cells were manually annotated, after which cells were detected using a watershed algorithm on the DAPI channel. To differentiate positive from negative cells, a Random Trees network classifier was trained in a semi-supervised way to detect positive cells based on signal intensity of either a combination of channels, or on a singular channel alone. Cell type abundance was expressed as number of positive detections normalized to total area.

### Spatial transcriptomics analysis

Visium data from 14 sarcoidosis patients skin biopsies^47^ was concatenated into one object within the scverse toolkit^68^. Raw counts were normalized, and log transformed. A set of 21 literature-derived genes were used to identify neutrophils: FCGR3B, FUT4, MPO, CSF3R, NAMPT, CD53, CAMP, CLEC7A, GCA, MNDA, FPR1, IL1R2, CTSG, ELANE, FCN1, LCN2, NAMPT, CD177, HCAR3, CMTM2, PROK2. The score genes function of Scanpy^69^ was used to calculate a relative expression score for each spot. Spots within the top 10% of the score distribution per sample and expressing at least 5 of the genes were marked as neutrophil-enriched. Spatial location of neutrophil-enriched spots and expression of genes of interest in both spot groups was visualized using Scanpy functionality. The relative expression of functional genes of interest (EGR2, IL4R, MRC1, PIM1) was quantified using the score genes function. Difference in gene score or normalized gene expression distributions was tested using a Wilcoxon signed-rank test with Benjamini-Hochberg correction for multiple testing.

### Analysis of publicly available gene expression data sets

Data from published array expression (from EMBL-EBI ; E-MTAB-13674)^49^ and RNA sequencing (NCBI GEO, GSE150996)^48^ was analyzed with regards to human alternative polarization markers.

### Statistics

Statistical analysis was performed using GraphPad Prism Version 10 (GraphPad, La Jolla). The detailed statistical tested are outlined in the respective figure legends. Depending on the group size two-tailed t-test or ANOVA was used. A *p*-value < 0.05 was considered statistically significant.

---

**Supplementary Figure 1:**
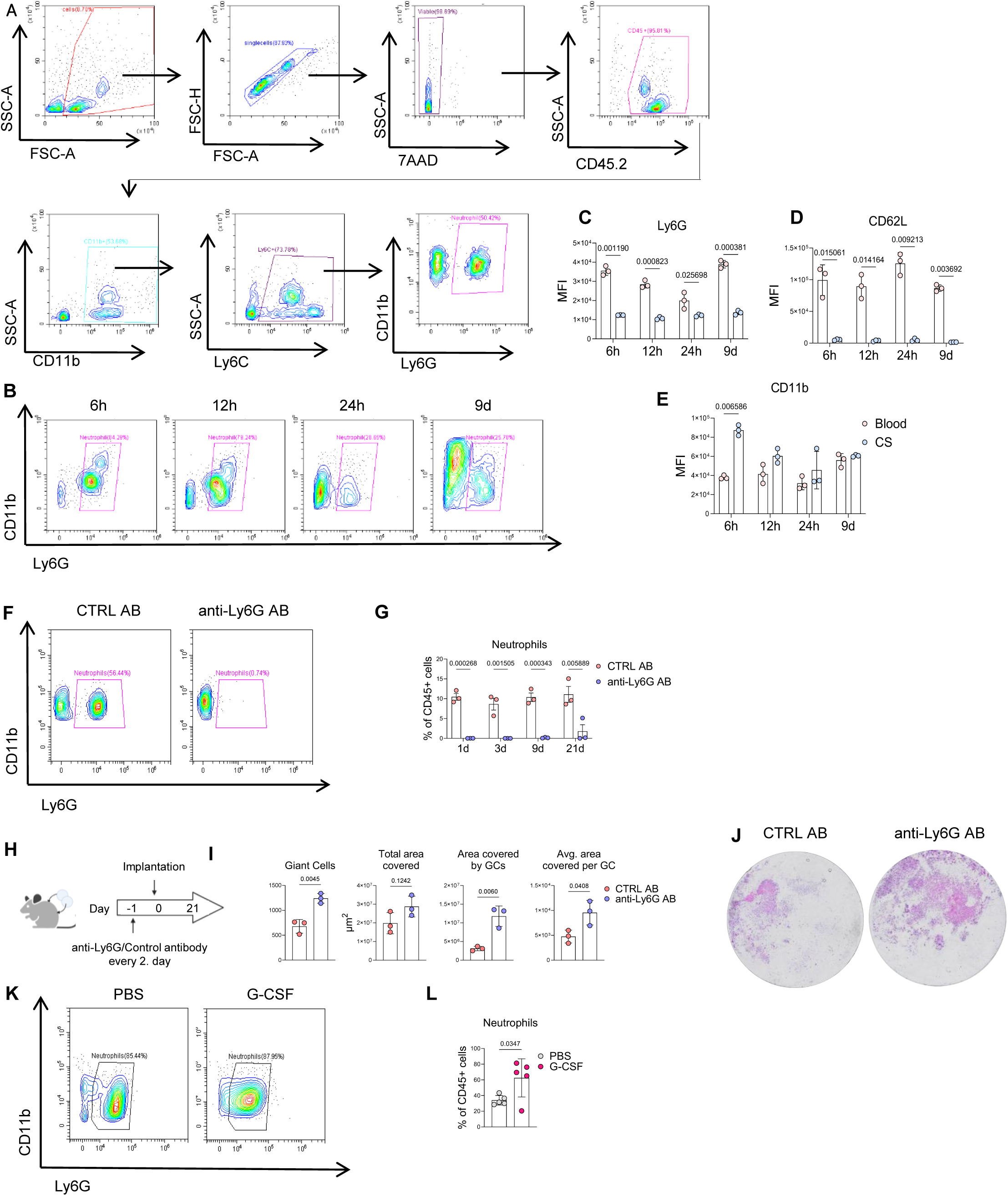
**(A)** Gating strategy for neutrophils used for cells isolated from blood and coverslips. **(B)** Representative flow cytometry plots of neutrophil population from cells isolated from coverslips 6h, 12h, 24h, and 9d after implantation. Neutrophils were gated as CD45+, CD11b+, Ly6C+, Ly6G+ cells. **(C-E)** Quantification of GeoMFI of Ly6G **(C),** CD62L **(D),** and CD11b **(E)** on neutrophils at different time points after coverslip implantation in blood and coverslip (n = 3 per time point). **(F)** Representative flow cytometry plots of neutrophils from blood from mice receiving either isotype control or anti-Ly6G depletion antibody 9 days after coverslip implantation. **(G)** Quantification of neutrophils in peripheral blood as percentage of CD45+ cells in mice receiving control or anti-Ly6G depletion antibody 1d, 3d, 9d, and 21d after implantation (n = 3 per time point). **(H)** Experimental outline of neutrophil depletion in mice upon coverslip implantation via anti-Ly6G depletion antibody for 21d. **(I)** Quantification of granulomatous burden upon anti-Ly6G depletion 21d after implantation. **(J)** Representative images of hematoxylin & eosin (H&E) staining of coverslips of mice either receiving control or anti-Ly6G depletion antibody 21 days after implantation. **(K)** Representative flow cytometry plots of neutrophils from blood from mice receiving either PBS or G-CSF 9 days after coverslip implantation. **(L)** Quantification of neutrophils in peripheral blood as percentage of CD45+ cells in mice receiving either PBS or G-CSF 9 days after coverslip implantation (n = 5 per condition). **(C-E, I, L)** Data are shown as mean ± standard deviation (SD) or **(G)** as mean ± standard error of the mean (SEM). Statistical analysis: multiple paired t tests **(C-E);** multiple unpaired t tests **(G)**; unpaired t tests **(I, L).**

**Supplementary Figure 2:**
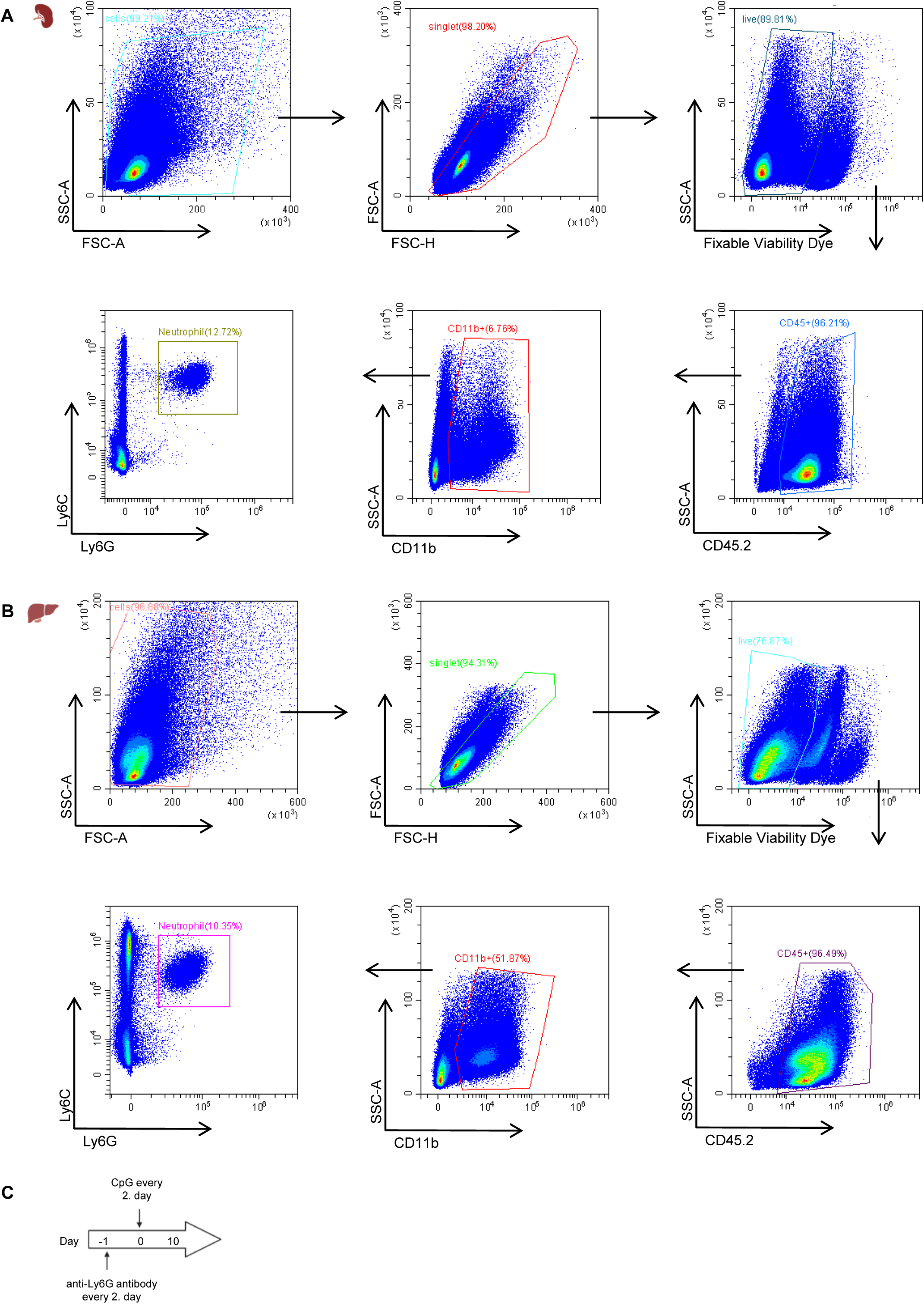
Gating scheme for neutrophils in spleen **(A)** and liver **(B)** in CpG induced murine MAS model as used in Figure 1O-P. **(C)** Schematic of neutrophil depletion in CpG-dependent murine MAS model.

**Supplementary Figure 3:**
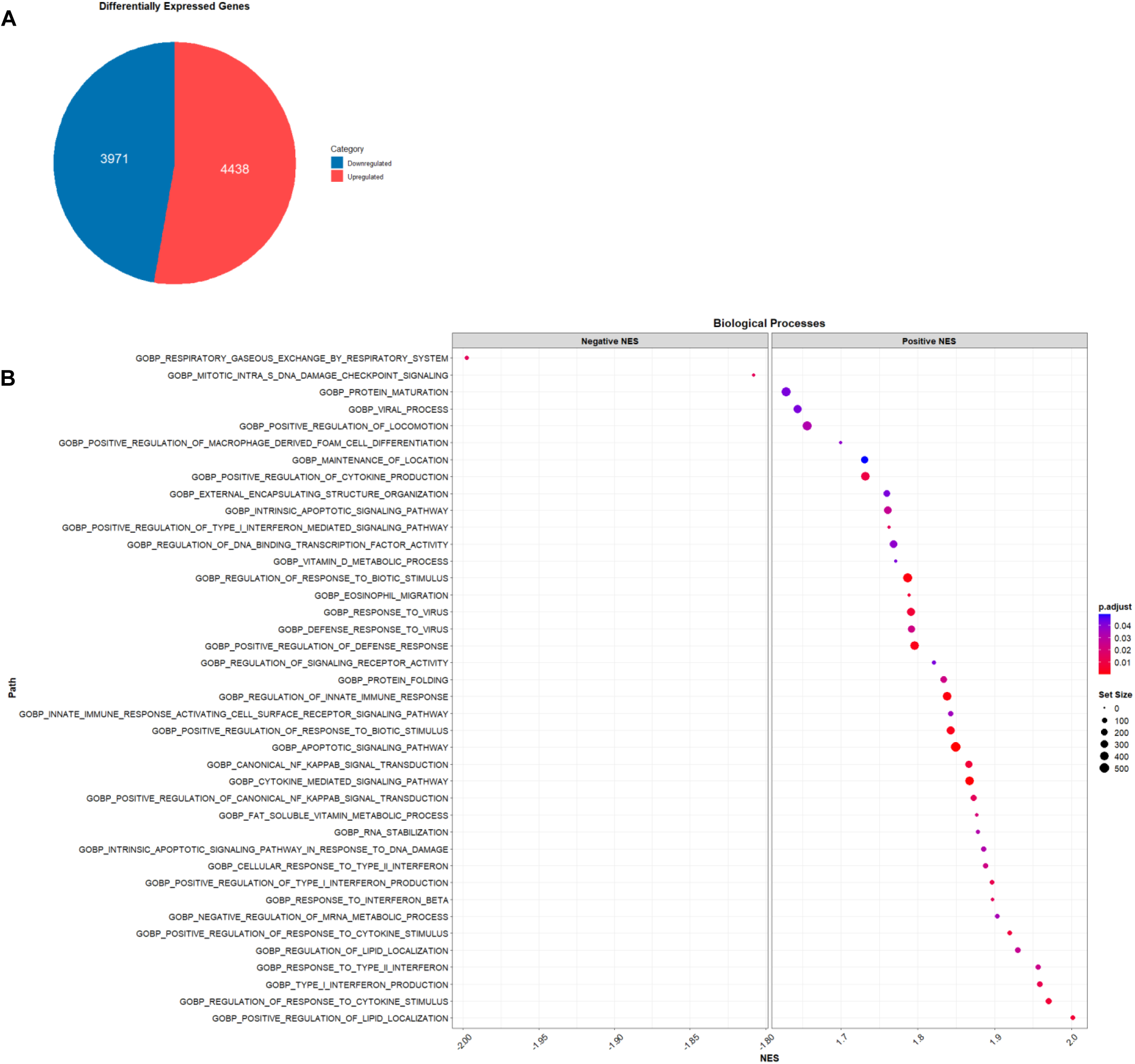
**(A)** Differentially expressed up- and down regulated genes in bulk RNA sequencing data set in CS vs. blood neutrophils. **(B)** Biological processes analysis in CS vs. blood neutrophils.

**Supplementary Figure 4:**
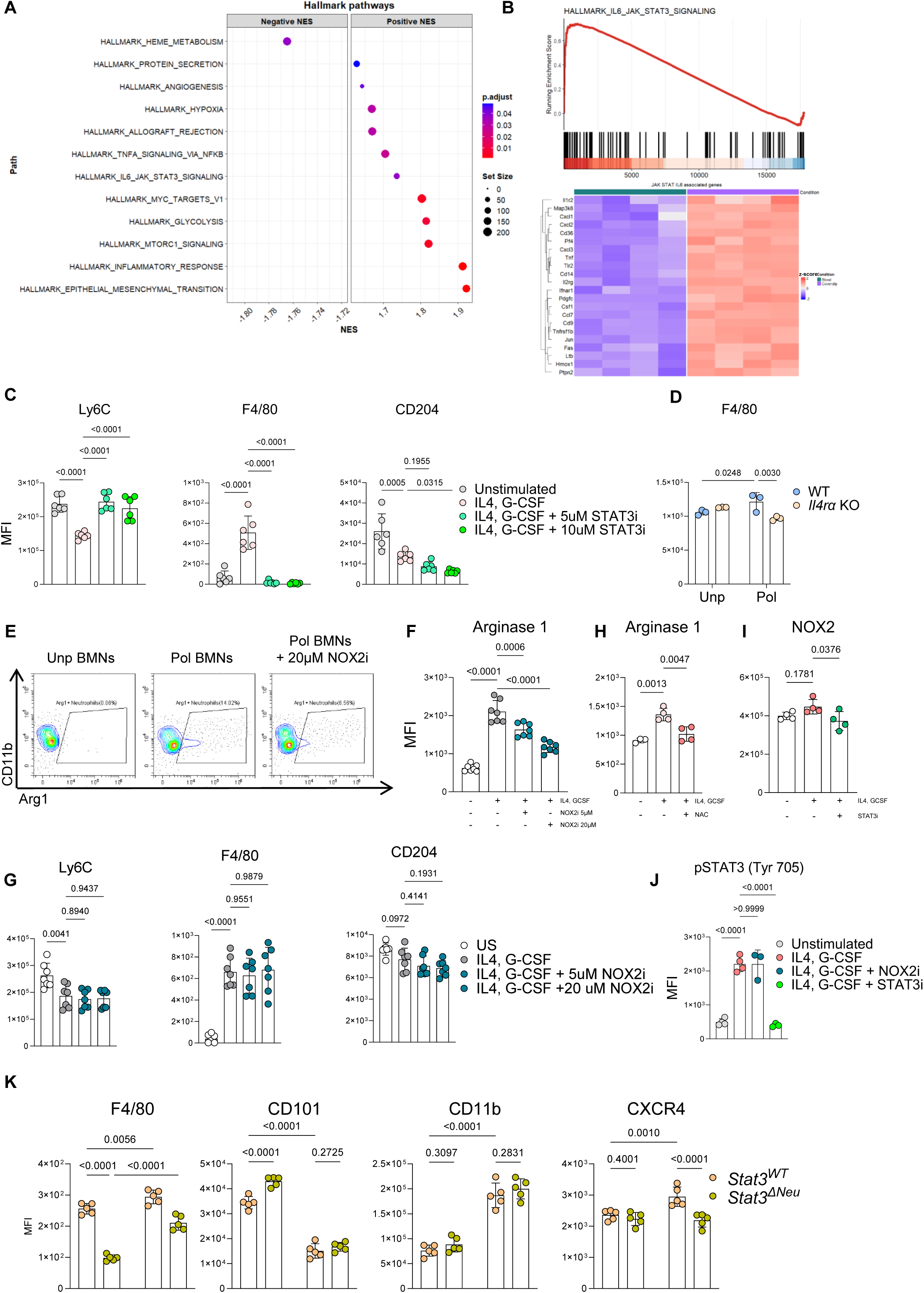
**(A)** Hallmark pathway analysis from bulk RNA sequencing data set. **(B)** GSEA and heat map of differentially expressed genes from IL6-JAK-STAT3 signaling pathway in blood vs. coverslip neutrophils. **(C)** Quantification of Ly6C, F4/80, and CD204 in Unp BMNs, Pol BMNs, and Pol BMNs treated with an STAT3 inhibitor for 24h, corresponding to Figure 3E, F. **(D)** Quantification of F4/80 in WT and Il4r*a* KO neutrophils upon alternative polarization, corresponding to Figure 3G - I. **(E)** Representative flow plots and **(F)** quantification of arginase 1 expression 24h after stimulation in Unp BMNs, Pol BMNs, and Pol BMNs treated with a NOX2 inhibitor (NOX2i) (n = 6). **(G)** Quantification of Ly6C, F4/80, and CD204 US BMNs, Pol MNs, and Pol BMNs treated with a NOX2 inhibitor for 24h, corresponding to Figure S4E - F. **(H)** Flow-cytometry based quantification of arginase 1 in Unp BMNs, and Pol BMNs treated with the ROS scavenger N-acetyl-cysteine (NAC) upon stimulation and treatment for 24h (n = 4). **(I)** Quantification of NOX2 expression in Unp BMNs, Pol BMNs, and Pol BMNs treated with a STAT3 inhibitor (n = 4). **(J)** Quantification of STAT3 phosphorylation in unstimulated neutrophils and upon cytokine stimulation in combination with a NOX2 or STAT3 inhibitor (n = 3). **(K)** Flow cytometric quantification of the GeoMFI of F4/80, CD101, CD11b, and CXCR4 in neutrophils in blood and coverslip from *Stat3^WT^* and *Stat3^ΔNeu^* mice 9 days after CS implantation, corresponding to Figure 3 M - P. **(C, F, G – J)** Data is depicted as mean ± standard deviation (SD). Data was analyzed using ordinary one-way ANOVA **(D, K)** or 2-way ANOVA. **(C, F – I).**

**Supplementary Figure 5:**
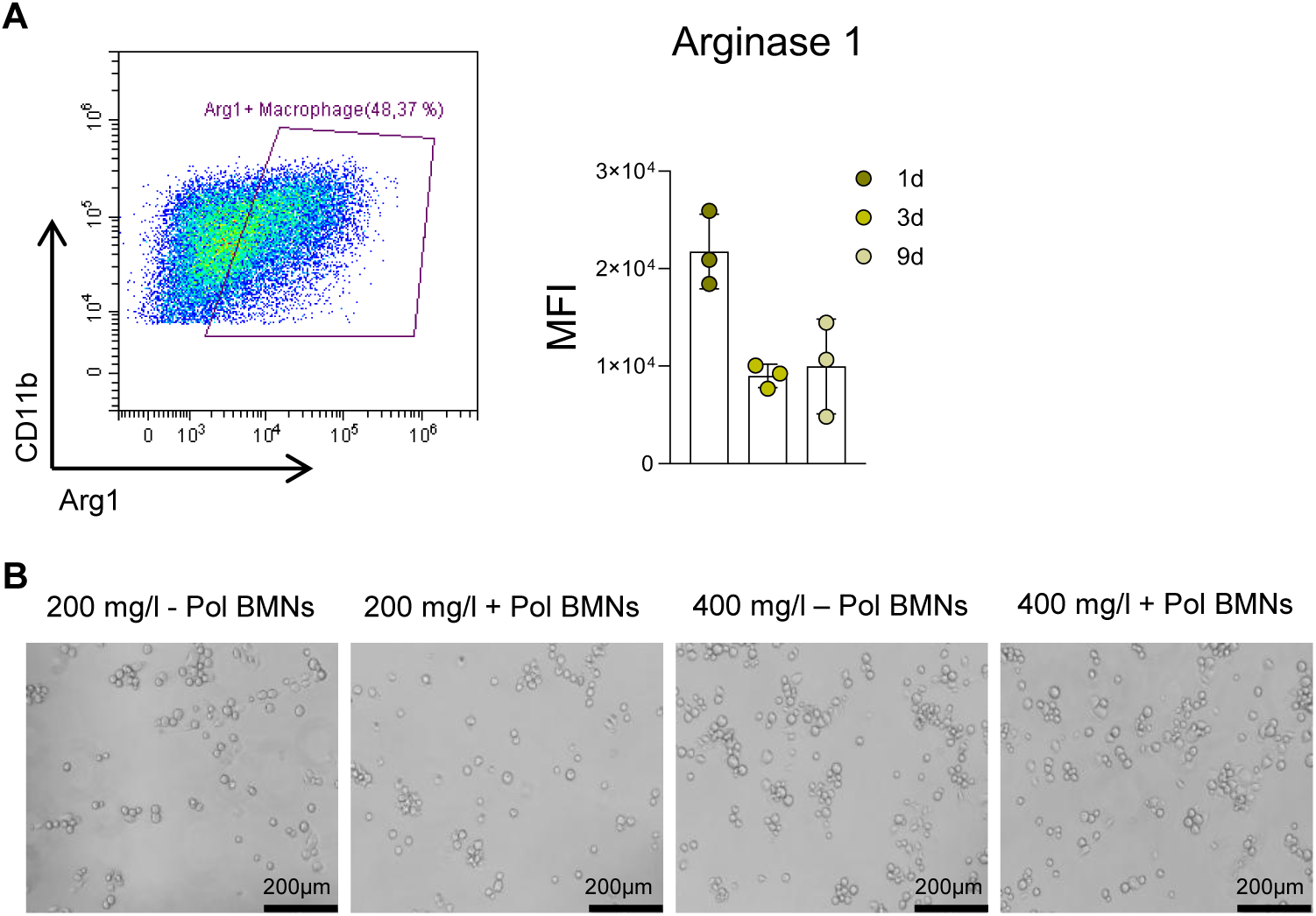
**(A)** Representative flow cytometry plot of arginase 1+ macrophages on the coverslip granuloma 3d after implantation and quantification of the MFI of arginase 1 in macrophages 1d, 3d, and 9d after coverslip implantation (n = 3 per condition). **(B)** Exemplary images of *Tsc2^ΔLyz2^* macrophage and neutrophil co-culture system quantified in Figure 4G, H. **(A)** Data is depicted as mean ± standard deviation (SD).

**Supplementary Figure 6:**
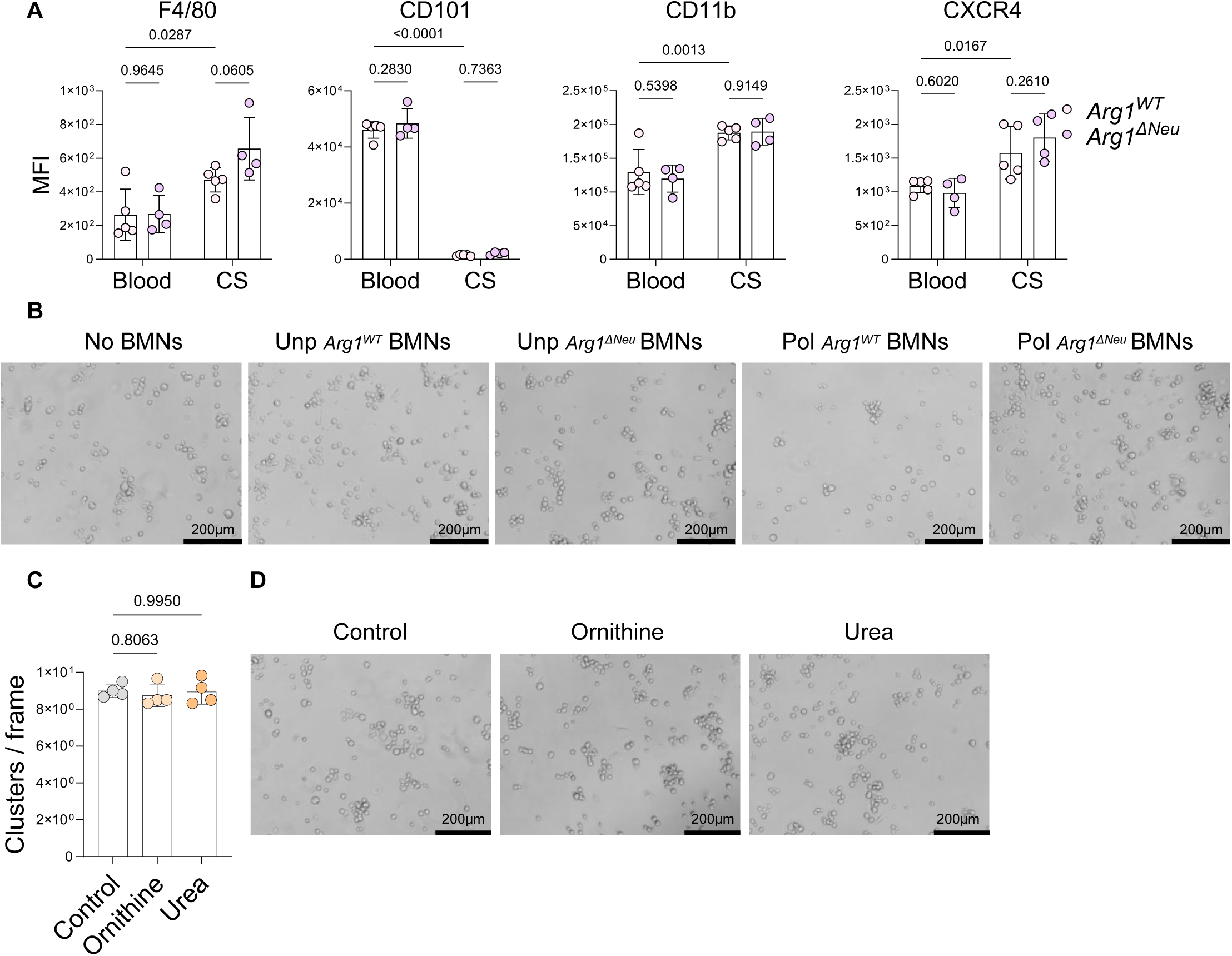
**(A)** Flow cytometric quantification of the GeoMFI of F4/80, CD101, CD11b, and CXCR4 in neutrophils in blood and coverslip from *Arg1^WT^* and *Arg1^ΔNeu^* mice 9 days after CS implantation, corresponding to Figure 5C. **(B)** Representative images of *Tsc2^ΔLyz2^* macrophage and neutrophil co-culture system quantified in Figure 5F. **(C)** Quantification and **(D)** representative images of *Tsc2^ΔLyz2^* macrophages cultured in with ornithine or urea compared to control (n = 4 per condition). Data are shown as means ± SD. Statistical analysis: RM-one-way ANOVA **(A, C)**.

**Supplementary Figure 7:**
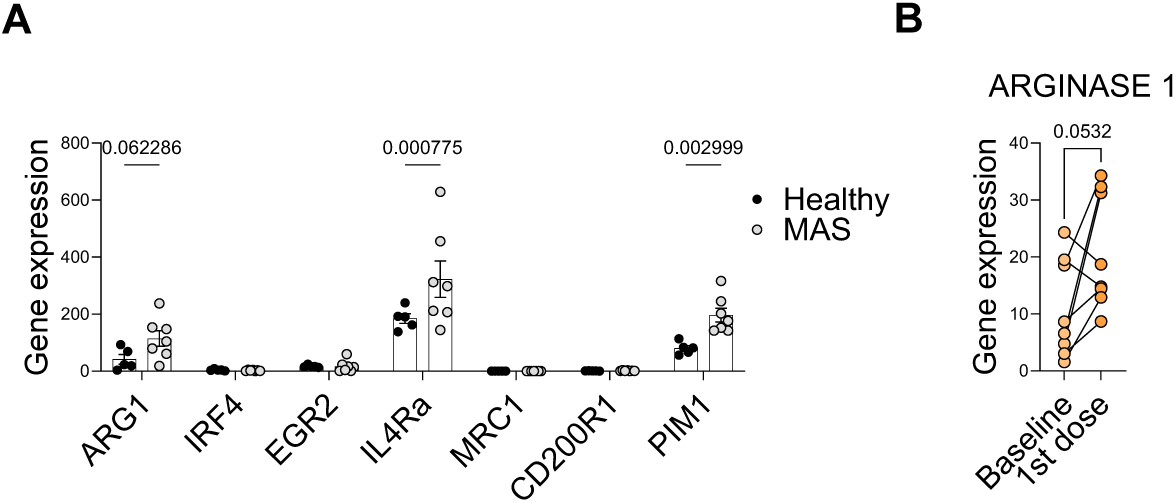
**(A)** Expression of human polarization markers in healthy vs. MAS patients (n = 5 - 7). **(B)** Gene expression of arginase 1 in CGD patients upon treatment with interferon-γ (n = 8). (**A, B**) Data are shown as means ± SD. Statistical analysis: multiple unpaired t-tests (**A**), paired t-test (**B**).

## Notes

### Competing Interest Statement

The authors have declared no competing interest.

